# SPACEc: A Streamlined, Interactive Python Workflow for Multiplexed Image Processing and Analysis

**DOI:** 10.1101/2024.06.29.601349

**Authors:** Yuqi Tan, Tim N. Kempchen, Martin Becker, Maximilian Haist, Dorien Feyaerts, Yang Xiao, Graham Su, Andrew J. Rech, Rong Fan, John W. Hickey, Garry P. Nolan

## Abstract

Multiplexed imaging technologies provide insights into complex tissue architectures. However, challenges arise due to software fragmentation with cumbersome data handoffs, inefficiencies in processing large images (8 to 40 gigabytes per image), and limited spatial analysis capabilities. To efficiently analyze multiplexed imaging data, we developed SPACEc, a scalable end-to-end Python solution, that handles image extraction, cell segmentation, and data preprocessing and incorporates machine-learning-enabled, multi-scaled, spatial analysis, operated through a user-friendly and interactive interface.

## Main

Multiplexed imaging allows simultaneous measurement of dozens to thousands of molecular markers (RNA, protein, or metabolome) within single cells with high resolution spatial positioning (1–4). Technologies such as co-detection by indexing (CODEX), multiplexed ion beam imaging (MIBI), and others that employ multiplexed antibody detection (5–11) provide incredibly detailed spatial data due to standardization of reagents and procedures (12–15). Standardization of techniques and methods has enabled the collection of amply detailed spatial data (16–21) that have revealed spatial relationships between cells, with appropriate analysis regimens, furthered our understanding of disease states (22, 23), tissue organization (15), and progression of cancer (24, 25). The meaning associated with such relationships depends on ready access to advanced analysis suites that are easy to use. Broad adoption of multiplexed imaging has been hindered by the absence of end-to-end, computationally efficient workflows and the need to integrate diverse codebases, environments, and algorithms. Most analysis frameworks focus either on preprocessing (e.g., Steinbock) or specific aspects of spatial data analysis (e.g., Spectre, SPIAT, Giotto, imcRtools) (26–29) (Supplementary Data Table 1-3). Some platforms offer more integrated solutions but require that users switch between tools and programming languages, posing difficulties for those with limited computational expertise. Moreover, some spatial-omics tools support both sequencing and imaging-based approaches but offer limited spatially resolved analyses beyond neighborhood identification (e.g., Squidpy) (30).

**Table 1.** Timing for resource-intensive steps in the SPACEc pipeline.

| Step | Name | Function | Specification | Timing |
| --- | --- | --- | --- | --- |
| 2 | Downscale entire tissue image | sp.hf.downscale_tissue | 1 qptiff image (3.3 GB, 28800x23040 px, 29 channels, 147128 cells) | downscale_factor = 64, 3min 20.3s, (25 GB of Swap used) |
| 3 | Rename tissue (optional) | sp.tl.label_tissue | 1 qptiff image (3.3 GB, 28800x23040 px) | downscale_factor = 64, 0.2s |
| 4 | Extract individual labeled tissues into separate tiff stacks | sp.tl.save_labelled_tissue | 1 qptiff image (3.3 GB, 28800x23040 px) | 8min 25.3s, (35 GB of Swap used) |
| 5 | Visualize segmentation channel (optional) | sp.pl.segmentation_ch | 1 Tiff image (403 MB, 2613x2614px) | 1.3s |
| 6 | Run cell segmentation | sp.tl.cell_segmentation | 1 Tiff images (403 MB, 2613x2614px), Mesmer segmentation | 55.7s |
| 7 | Visually inspect segmentation results | sp.pl.show_masks | 1 Tiff image (403 MB, 2613x2614px) | 2.3s |
| 8 | Load the segmented results | sp.pp.read_segdf | 2 csv files (27.5 and 34.5MB) | 0.4s |
| 9 | Initial filtering of artifacts/debris by DAPI intensity and cell size | sp.pp.filter_data | 52376 cells, 71 features | 1.6s |
| 10 | Normalize data | sp.pp.format | 52376 cells, 66 features | 0.001s |
| 11 | Second filtering for noisy cell | sp.pl.zcount_thres<br>sp.pp.remove_noise | 52376 cells, 57 features | 0.4s |
| 12 | Cell type annotation via clustering | sp.tl.clustering | 52376 cells, 39 features, leiden clustering | 1min 0.8s |
| 13 | Visualize & annotate clustering results | sc.pl.umap<br>sc.pl.dotplot<br>sp.pl.catplot | 52376 cells, 57 features | 5s + time needed for manual annotation |
| 14 | Compute basic statistics of the cell type composition | sp.pl.stacked_bar_plot<br>sp.pl.create_pie_charts | 52376 cells, 57 features | 0.3s |
| 15 | Cell type annotation via machine-learning classification | sp.tl.ml_train<br>sp.tl.ml_predict | 52376 cells, 57 features | 13.6s |
| 16 | Prepare data for an interactive session | sp.tl.tm_viewer | 52376 cells, 17 cell types | 2.9s |
| 17 | Additional interaction data exploration (optional) |  |  |  |
| 18 | Compute cellular neighborhoods | sp.tl.neighborhood_analysis | 52376 cells, 17 cell types | 0.4s |
| 19 | Visualize & annotate the cellular neighborhoods analysis results | sp.pl.cn_exp_heatmap<br>sp.pl.catplot | 52376 cells, 17 cell types | 1.5s generation of figures + time needed for manual annotation |
| 20 | Generate spatial context maps | sp.tl.cn_map | 52376 cells, 17 cell types | 0.6s |
| 21 | Create cellular neighborhood interface analysis via barycentric coordinate plot | sc.pl.bc_project | 52376 cells, 5 CNs | 2s |
| 22 | Compute for patch proximity analysis | sp.tl.patch_proximity_analysis | 52376 cells, 5 CNs | 29.1s |
| 23 | Calculate cell-cell interaction | sp.tl.identify_interactions<br>sp.tl.filter_interactions<br>sp.pl.dumbbell | 52376 cells, 17 cell types, 100 iterations | 49s |

The Structured Spatial Analysis for Codex Exploration (SPACEc) framework was designed to enable researchers to unlock the full potential of these multiplexed imaging data using an interactive Python-based analysis pipeline. SPACEc performs image extraction, cell segmentation, single-cell data preprocessing and normalization, interactive quality inspection and annotation, and single-cell spatial analysis. SPACEc was designed with CODEX data in mind, but the workflow can now be extended to analyze data from platforms such MIBI and IMC, allowing for the modular introduction of other analysis approaches.

SPACEc provides solutions for accelerating cell-type annotation on large datasets through GPU-accelerated unsupervised clustering and supervised machine learning (ML) techniques. As such, the computational pipeline can efficiently process large image sizes (ranging from 8 to 40 gigabytes per image). Among the ML methods included for cell type annotations are support vector machines (SVM) and STELLAR, a geometric deep learning approach. SPACEc can be used on consumer and commercial GPUs alike for further speedups. SPACEc operates in Linux, MacOS, and Windows environments. We also provide Docker containers to simplify installation. To assist in implementation, a comprehensive step-by-step guide is provided on GitHub (https://github.com/yuqiyuqitan/SPACEc).

SPACEc incorporates the following steps for analyzing multiplexed images (**Fig. 1**): tissue extraction, cell segmentation and visualization, data preprocessing and normalization, cell-type annotation, interactive data inspection and exploration, and spatial analysis. To perform these steps, SPACEc integrates analytical tools that were developed specifically for this pipeline and tools that were previously established (20, 22, 31, 32). To identify underlying cellular biology within the terabytes of raw imaging data, SPACEc includes two cell segmentation algorithms, Mesmer (33) and Cellpose (34), developed by others, and quality assurance measures with visual feedback to ensure data accuracy. Once single-cell data is segmented and labeled, established standardized processes for normalization are employed (35). For interactive visualization, SPACEc employs TissUUmaps, a browser-based, GPU-accelerated tool (36). TissUUmaps is capable of visualizing 10^7^ data points simultaneously; general-purpose viewers like Napari, matplotlib, BigDataViewer, and QuPath cannot visualize millions of data points simultaneously due to CPU and RAM constraints (36–40). TissUUmaps supports the sharing of visualizations, has HDF5 and Anndata compatibility, and facilitates explorative analysis via either a web interface or a standalone application. SPACEc is compatible with existing analysis packages within the scverse ecosystem and can be employed to analyze data from recently developed frameworks for single-cell analysis (30, 41–43). The SPACEc toolkit also includes clustering options, including Leiden, Louvain, and FlowSOM, for identifying different groups of cell types (44–46). Cell-type labels can be transferred across datasets using an SVM (47). Furthermore, the SPACEc pipeline includes interactive modules to ensure data quality and to allow inspection of cell-type labels in their original tissue coordinates. SPACEc offers five advanced spatial analysis algorithms; detailed documentation for these analyses can be found in the (**Supp. Note 1** and **Supp. Note 2**) section.

**Figure 1:**
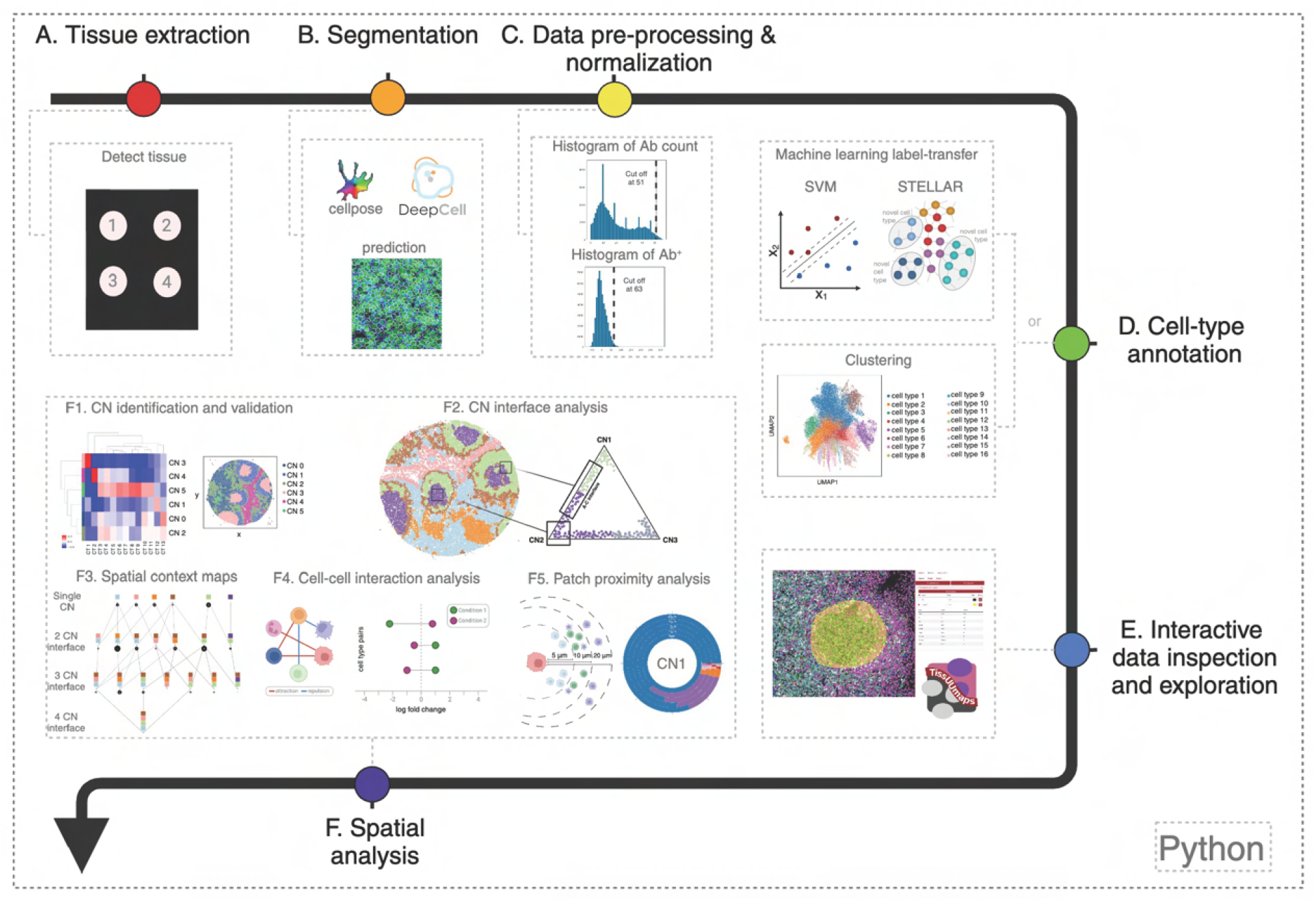
The SPACEc workflow. A) Data for individual tissue images are extracted from the qptiff file into separate tif files. B) Image segmentation is performed on individual tif files. C) The segmented data frame with quantified marker intensities is filtered, normalized, and saved as an Anndata object. D) Each cell in the processed Anndata object is annotated through either unsupervised clustering or machine-learning-based annotation. E) The annotation results are inspected and annotated interactively using TissUUmaps. F) Spatial analysis, including CN analysis, spatial context mapping, CN interface analysis, patch proximity analysis, and cell-cell interaction analysis, is performed. Results are stored within the Anndata object. The pipeline can be customized to integrate with other tools.

To illustrate the utility of SPACEc, we processed CODEX Phenocycler datasets from healthy and inflamed tonsils. First, the pipeline extracted and labeled individual tonsil tissues from the tissue microarray (**Fig. 2A**). Mesmer was used to segment the data, the resultant segmentation masks were visualized (**Supp. Fig. 1A**), data were filtered and normalized (**Supp. Fig. 1B**), and then stored in an Anndata format, all within SPACEc. Next, cell types were annotated with Leiden clustering and with cell-type-specific markers (**Fig. 2B** and **Supp. Fig. 2A**). The cell-type labels were then mapped back to their original spatial coordinates to verify the accuracy of cell-type labels in the context of known biology (**Fig. 2B**). The cell-type compositions can be visualized as a pie chart or stacked bar chart (**Supp. Fig. 2B-D**). We also tested the accuracy of annotating cell types with SVM, leveraging annotated healthy tonsil data as training and unannotated inflamed tonsil as test data and observed similar performance (**Supp. Fig. 2E**). Finally, we used TissUUmaps (40) to interactively inspect the labels and conduct preliminary analysis (**Fig. 2D** and **Supp. Fig. 3**).

**Figure 2:**
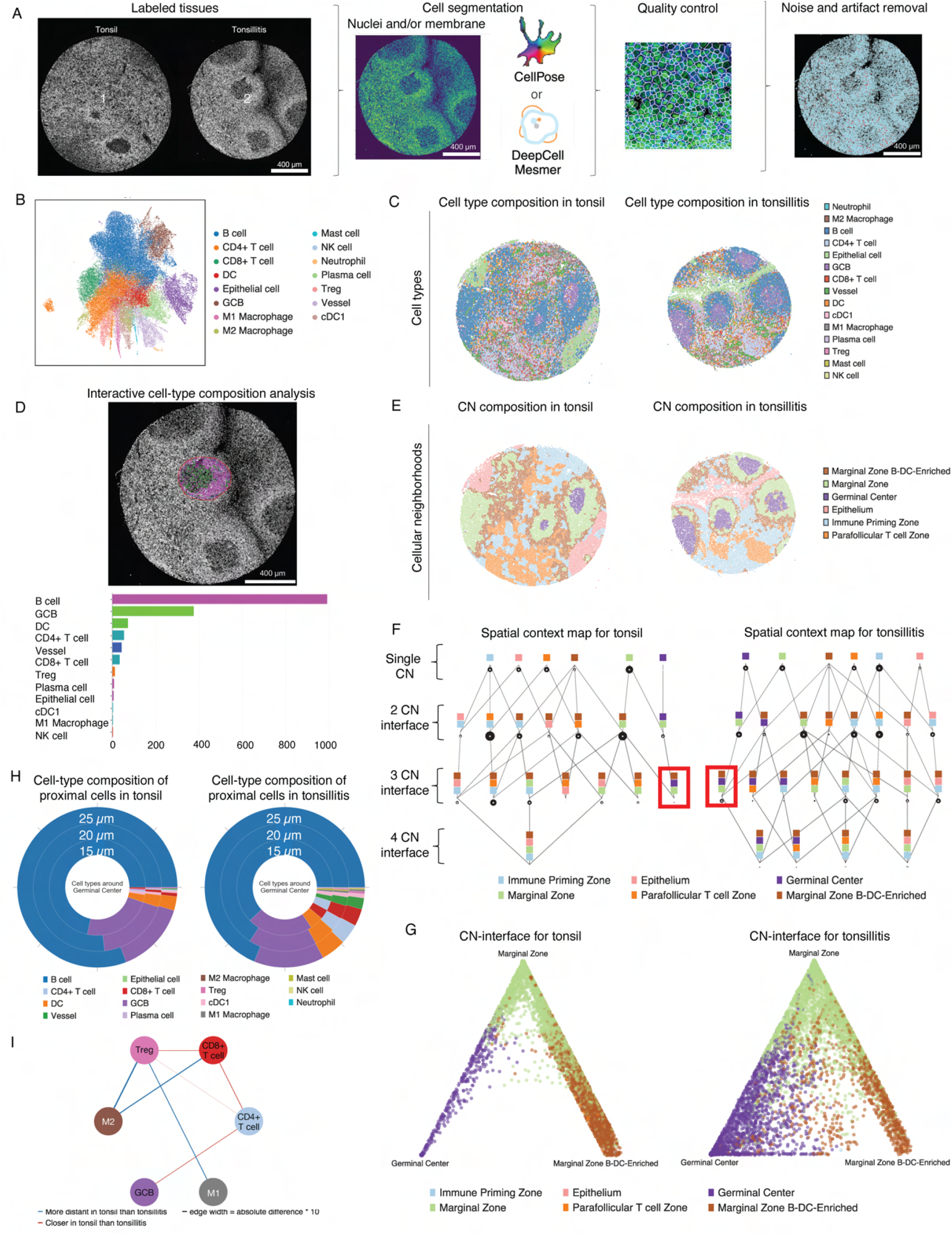
An illustration of SPACEc’s workflow with tonsil Phenocycler data. A) In the example, there are two CODEX-imaged tissue cores from a larger tissue microarray (tonsil and tonsillitis). The two tissues were detected, labeled, extracted, segmented, and visualized individually by SPACEc. B) UMAP visualization of annotated cells. C) Cells from healthy (left) and inflamed (right) tonsil are visualized on original tissue coordinates. Each dot represents a cell. D) Region selection using the Plot Histogram plug-in (top) enables the display of cell-type composition within that region as a histogram (bottom). E) CNs from healthy (left) and inflamed (right) tonsil visualized on original tissue coordinates. Each dot represents a cell. F) Spatial context maps of healthy (left) and inflamed (right) tonsil. Rows show the number of neighborhoods in combinations: row 1, a single neighborhood accounts for at least 85% of the neighborhoods surrounding the window; row 2, two neighborhoods make up more than 85% of the neighborhoods in the window; and row 3 and beyond, multiple neighborhoods are present in the window. G) Barycentric plots of window combinations in healthy (left) and inflamed (right) tonsil. H) Percentages of cell types that surround all patches in germinal center CN in healthy (left) and inflamed (right) tonsil. I) Cell-cell interactions in healthy (left) and inflamed (right) tonsil were visualized using distance network graphs. Blue edges indicate the cell-type distances that are further and red edges indicate distances that are closer in healthy than in inflamed tonsil. The edge width is proportional to the fold change between the healthy and inflamed samples.

To showcase the analytic capacity of SPACEc, we performed the advanced spatial analyses incorporated in the pipeline. Six cellular neighborhoods (CNs) were identified and annotated based on enriched cell types (**Fig. 2E** and **Supp. Fig. 4**). The inflamed tonsil has more mature germinal centers than the healthy tonsil, in accordance with the colonization and proliferation of B cells following initial activation of antigen-specific B cells. To inspect higher-level tissue architectural differences, spatial context maps for each tissue were computed (**Fig. 2F** and **Supp. Fig. 4**). Based on the hierarchical layout of spatial context maps, the interface between three CNs (Marginal Zone, Germinal Center, and Marginal Zone Enriched in B Cells and Dendritic Cells) was different between the two conditions. In a barycentric coordinate plot, we observed that the interface between the Germinal Center CN and the Marginal Zone Enriched in B Cells and Dendritic Cells CN was particularly pronounced in the inflamed tonsil (**Fig. 2G** and **Supp. Fig. 6**), suggestive of an efficient antigen response and memory B cell formation.

In addition to delineating the intrinsic tissue architecture, SPACEc can be used to inspect cell-cell and CN-CN interactions. For instance, the composition of cells surrounding a defined tissue structure, such as the germinal center CN can be calculated. We projected concentric circles radiating in distances from 15-25 µm from the border cells and observed an enrichment of CD8^+^ T cells and CD4^+^ T cells surrounding the Germinal Center CN in the inflamed tonsil (**Fig. 2H** and **Supp. Fig. 7**). Similar patch proximity analysis can be extended to analyze the CNs composition bordering each CN. Lastly, we looked at the changes in cell-cell interactions between the two conditions (**Figure 2I** and **Supp. Fig. 8**). To evaluate differences in cell-cell interactions between the two conditions, the distance between pairs of cell types within a threshold (i.e., 100 µm) is calculated, and then cell-type labels are randomly permuted within each tissue section to simulate random chance conditions. In tonsil samples, CD8^+^ T cells are closer to dendritic cells in the inflamed tonsil than the healthy tonsil, indicative of T cell priming and activation during tonsillitis.

Here we showed that SPACEc provides a comprehensive and structured Python-based workflow for analysis of multiplexed images. The pipeline performs essential steps of tissue extraction, cell segmentation, and visualization, data preprocessing and normalization, as well as cell-type annotation. Furthermore, it enables interactive data inspection and spatial analysis with outputs in various formats. SPACEc incorporates a range of analytical tools to ensure a thorough and versatile analysis process. With detailed step-by-step instructions, users of all levels can implement the workflow, efficiently navigating spatial landscapes and extracting valuable biological insights from highly multiplexed images.

## Methods

### Tissue samples

Tonsil cores were extracted from a larger multi-Tumor tissue microarray (TMA), which included a total of 66 unique tissues (51 malignant and semi-malignant tissues, as well as 15 non-malignant tissues). Representative tissue regions were annotated on corresponding hematoxylin and eosin (H&E)-stained sections by a board-certified surgical pathologist (S.Z.). Annotations were used to generate the 66 cores each with cores of 1mm diameter. FFPE tissue blocks were retrieved from the tissue archives of the Institute of Pathology, University Medical Center Mainz, Germany, and the Department of Dermatology, University Medical Center Mainz, Germany. The multi-tumor-TMA block was sectioned at 3µm thickness onto SuperFrost Plus microscopy slides before being processed for CODEX multiplex imaging as previously described (13).

### CODEX multiplexed imaging and processing

To run the CODEX machine, the slide was taken from the storage buffer and placed in PBS for 10 minutes to equilibrate. After drying the PBS with a tissue, a flow cell was sealed onto the tissue slide. The assembled slide and flow cell were then placed in a PhenoCycler Buffer made from 10X PhenoCycler Buffer & Additive for at least 10 minutes before starting the experiment. A 96-well reporter plate was prepared with each reporter corresponding to the correct barcoded antibody for each cycle, with up to 3 reporters per cycle per well. The fluorescence reporters were mixed with 1X PhenoCycler Buffer, Additive, nuclear-staining reagent, and assay reagent according to the manufacturer’s instructions. With the reporter plate and assembled slide and flow cell placed into the CODEX machine, the automated multiplexed imaging experiment was initiated. Each imaging cycle included steps for reporter binding, imaging of three fluorescent channels, and reporter stripping to prepare for the next cycle and set of markers. This was repeated until all markers were imaged. After the experiment, a .qptiff image file containing individual antibody channels and the DAPI channel was obtained. Image stitching, drift compensation, deconvolution, and cycle concatenation are performed within the Akoya PhenoCycler software. The raw imaging data output (tiff, 377.442nm per pixel for 20x CODEX) is first examined with QuPath software (https://qupath.github.io/) for inspection of staining quality. Any markers that produce unexpected patterns or low signal-to-noise ratios should be excluded from the ensuing analysis. The qptiff files must be converted into tiff files for input into SPACEc. Data preprocessing includes image stitching, drift compensation, deconvolution, and cycle concatenation performed using the Akoya Phenocycler software. The raw imaging data (qptiff, 377.442 nm/pixel for 20x CODEX) files from the Akoya PhenoCycler technology were first examined with QuPath software (https://qupath.github.io/) to inspect staining qualities. Markers with untenable patterns or low signal-to-noise ratios were excluded from further analysis. A custom CODEX analysis pipeline was used to process all acquired CODEX data (scripts available upon request). The qptiff files were converted into tiff files for tissue detection (watershed algorithm) and cell segmentation.

### Cell segmentation

SPACEc includes two segmentation methods: Deepcell/Mesmer (33) and Cellpose (34). SPACEc uses the pre-trained multiplexed imaging model for Mesmer and allows access to the model zoo of Cellpose. Additionally, users can train their own Cellpose models and incorporate them into the pipeline. Users can input multiple channels for segmentation. Both Mesmer and Cellpose methods require the nuclei channel as minimum input and allow input of additional markers (for example, HLA-ABC for marking membranes). After segmentation, mean intensities are quantified for each channel in every mask. In this manuscript, we used Mesmer for the cell segmentation.

### Normalization

*Z Normalization*: Individual marker intensities were Z-normalized for all cells within the dataset. This step aimed to standardize the range of each marker, considering variations in fluorescent intensities due to antibody staining strength and exposure times. *Log (Double Z) Normalization*: Initially, Z normalization was conducted on each marker intensity, followed by another Z normalization applied to each cell. These values were subsequently transformed into probabilities. Finally, a negative log transformation was applied to the complement of the probabilities. By equalizing signal intensities through the first Z normalization, a comparison of marker Z scores became feasible. Moreover, the second Z normalization helped identify positive markers with high probability among cells, which typically exhibit positivity for only a subset of the numerous markers recognized by antibodies. The negative log transformation of the complement of probability amplifies values with high probabilities, facilitating their utilization in clustering algorithms. *Min_Max Normalization*: Initially, the 1st and 99th percentiles were determined to set the minimum and maximum values, respectively, for each fluorescent channel. Subsequently, each value in the channel underwent normalization by calculating the difference between the minimum value and the range of values. Capping values at the 99th percentile assists in eliminating artificially high background fluorescent intensities often observed in imaging datasets. *Arcsinh Normalization*: This technique involved an arcsinh transformation on marker intensities, followed by scaling the resulting values with a cofactor of 150. Such normalization is suitable for datasets containing low or negative values resulting from background subtraction. In this study, we used z-normalization.

### Single-cell matrix preprocessing and cell clustering

Very small cells and cells with low DNA marker expression should be excluded from the analysis before z-score normalization and filtering of the cells as previously described (35). After pre-processing the single-cell matrix cells are clustered using Leiden clustering as implemented in the Scanpy library. For example, cells in the tonsil dataset were clustered with a resolution of 0.4 and 10 nearest neighbors. Subsequently, clusters are plotted in a heatmap and manually annotated based on the marker expression.

### SVM-based annotation of cell types

The SVM function is implemented in the standard scikit-learn SVM. Users must provide training data with labels. SVMs determine a decision boundary that separates different types of cells based on input protein features. The trained SVM model is then applied to a new dataset, which ideally has overlapping features, to transfer the cell-type labels.

### Cellular neighborhood analysis

Users first will select a window size (e.g. 20) of nearest neighbors. Within each window, we quantified the cells of each type, generating vectors representing cell counts. Subsequently, these vectors were subjected to clustering to identify commonly composed neighborhoods. After an extensive parameter search, users can identify the optimal number of cellular neighborhoods that provide unique insights into the spatial organization and heterogeneity of cell types within the tissue. Then a heatmap can be used to identify the cell types uniquely enriched in each neighborhood, which can be used for annotations. Six cellular neighborhoods were selected based on the distinctness of their enriched cell types.

### Spatial context map

First, a large (e.g. 70) window size of nearest neighbors to create the composition vectors. Within each window, we will identify the fewest neighborhoods that will make up more than 85 percent of the neighborhoods. This combination informs about prominent associations of neighborhoods in the window, a feature we term spatial context. Third, we counted each combination and connected the most prevalent combinations into a spatial context map. This hierarchical spatial context map shows different levels of neighborhood combinations and their relative frequencies.

### Cellular neighborhood interface analysis

To generate a barycentric coordinate projection, users need to select three CNs. Thereafter, large windows were applied to every cell (window size of 70 nearest neighbors). Window composition was analyzed in terms of the percentage distribution of CNs. Windows containing less than the user-defined threshold of 90 percent or a user-defined percentage of the three selected neighborhoods were excluded from the analysis.

### Patch proximity analysis

This method analyzes the composition of cells in the spatial proximity of patches. Patches are detected in spatial single-cell data by running HDBSCAN clustering over all centroids in a user-defined region that belongs to a previously specified CN. The resulting clusters are used as patches, and cells that are not part of a patch are ignored. The outermost cells of each patch are selected by constructing a concave hull and selecting the spanning points. To include cells close to the outlining border of the patch and to correct for cells that are distant from the patch, three nearest neighbors for each edge cell are included as well, resulting in a group of cells surrounding each patch. Subsequently, the selected cells are used as anchor cells to span a user-defined radius around each selected cell. For the tonsil data, the radius was X pixels. Cells within the radius that do not belong to the patch are counted as cells within spatial proximity. In this manuscript, we examined cells and CNs with the proximal distances at 15um, 20um, and 25um of the border.

### Cell-cell interaction analysis

Briefly, Delaunay triangulation for each cell is determined based on x, y positions within each field of view using the default settings from the SciPy package. To identify interacting cells and their coordinates, relevant information is extracted automatically from the output. Subsequently, the distances between cells connected by the edges of a Delaunay triangle in the two-dimensional space are computed using the following formula: 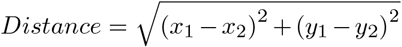. Cell-to-cell interactions within 265 pixels are identified. To establish a reference distribution of distances, 100 iterations of the triangulation calculation are conducted. In each iteration, the cells and their neighborhoods within each field of view are randomly reassigned to existing x, y positions. The average distances of cell-cell interactions for each field of view in each permutation are calculated and compared to the observed distances using a Mann-Whitney U Test. The fold enrichments of distances between the observed data compared to the mean distances derived from the permutation test are determined. The log fold changes of distance for each pair of interactions for which p values are less than 0.05 are plotted.

### Hardware

All procedures were tested on multiple platforms to ensure wide compatibility. Platforms were a workstation running Ubuntu 22.4 equipped with an AMD Threadripper 5955wx, 32 GB of RAM, and a Nvidia RTX A4500 GPU; a MacBook 2019 running MacOS 14.1.1 with a 2.6 GHz 6-Core Intel Core i7 and 16 GB of RAM; an Apple M1 12.4 MacBook Pro with 32 GB of RAM; and a Windows Server 2019 with an AMD Threadripper, 256 GB of RAM and dual RTX 3080.

## Code availability

All software and code used to produce the findings of this study, including all main and supplemental figures, are available at https://github.com/yuqiyuqitan/SPACEc.

## Data availability

A CODEX dataset of healthy and inflamed tonsil is used as an example to demonstrate the entire analytic workflow. The data were generated as part of a previous study using the PhenoCycler Fusion platform (6). The data can be accessed at [Dryad](https://datadryad.org/stash/share/OXTHu8fAybiINGD1S3tIVUIcUiG4nOsjjeWmrvJV-dQ) and contain the following files: 1) one example tiff file as input for tissue extraction, 2) channel markers in a txt file for the tiff file as input for cell segmentation, 3) an Anndata object that contains previously annotated data as reference for cell-type annotation using SVM-based annotation. An example of the directory for the folder that users will download to execute SPACEc functions is shown below.

1. Raw_data
  - tonsil_TMA.tif
  - channelnames.txt
2. Processed_data
  - adata_nn_demo_tonsil.h5ad

## ACKNOWLEDGEMENTS

This work was supported by the US National Institutes of Health (P01HL108797, U01AI101984, 5U54CA209971, 5U01AI140498, U54HG010426, U19AI100627, 5P01AI131374, UH3DK114937, U19AI135976, U2CCA233238, U2CCA233195, U19AI057229, U54HG012723); the US Food and Drug Administration (HHSF223201610018C, DSTL/AGR/00980/01); Cancer Research UK (C27165/A29073); the Bill and Melinda Gates Foundation (OPP1113682); the Cancer Research Institute; the Parker Institute for Cancer Immunotherapy (PICI0025); Hope Realized Medical Foundation (209477); the Kenneth Rainin Foundation (2020-1463); the Beckman Center for Molecular and Genetic Medicine; Celgene (133826, 134073); Vaxart, Inc. (202627); and the Rachford & Carlotta A. Harris Endowed Chair to G.P.N. Y.T. is supported by a Stanford Dean’s Fellowship and the Stanford Cancer Institute Cancer Innovation Award. J.W.H. was supported by an NIH T32 Fellowship (T32CA196585) and an American Cancer Society—Roaring Fork Valley Postdoctoral Fellowship (PF-20-032-01-CSM). M.H. is supported by the Deutsche Forschungsgemeinschaft (DFG, German Research Foundation) (project number: HA 9793/1-1). The work by M.B. was supported by the Federal Ministry of Education and Research (BMBF), Germany (01IS22077). D. F. is supported by a Society for Reproductive Investigation and Bayer Innovation/Discovery Grant and the Stanford Maternal and Child Health Research Institute Postdoctoral Support Award. This article reflects the views of the authors and should not be construed as representing the views or policies of the FDA, NIH, BMGF, or other institutions that provided funding. Some figures were created with BioRender.com.

## AUTHOR CONTRIBUTIONS

Conceptualization: Y.T., G.P.N.

Algorithm Development and Implementation: Y.T., T.N.K, M.B., J.W.H.

Analysis: Y.T., T.N.K, M.B., J.W.H.

Contribution of Key Reagents and technical expertise: M.H., D.F., Y.X., G.S., A.J.R.

Supervision: G.P.N.

Both Y.T. and T.N.K. contributed equally and have the right to list their name first in their CV.

## CONFLICT OF INTERESTS

G.P.N. received research grants from Pfizer, Inc.; Vaxart, Inc.; Celgene, Inc.; and Juno Therapeutics, Inc. G.P.N. has equity in Akoya Biosciences, Inc. G.P.N. is a scientific advisory board member of Akoya Biosciences, Inc.

## Supplementary Tables

**Supplementary Table 1:** The accessibility of essential analytical procedures for multiplexed image analysis within each computational package.

|  | Cell segmentation | Data pre-processing | Data normalization | ML cell type annotation | Downstream analysis | Interactivity |
| --- | --- | --- | --- | --- | --- | --- |
| **SPACEc** | Yes | Yes | Yes | Yes | Yes | Yes |
| Steinbock | Yes | No | No | No | No | Yes |
| imcRtools | No | Yes | Yes | Yes | Yes | No |
| SPIAT | No | No | No | No | Yes | No |
| Giotto | No | Yes | Yes | No | Yes | No* |
| Spectre | No | No | Yes | Yes | Yes | No |
| MCMICRO | Yes | Yes | Yes | No | Yes | Yes |

**Supplementary Table 2:** The programming languages employed for key analytical processes in multiplexed image analysis within each computational package.

|  | Cell segmentation | Data pre-processing | Data normalization | ML cell type annotation | Downstream analysis | Interactivity |
| --- | --- | --- | --- | --- | --- | --- |
| **SPACEc** | Python | Python | Python | Python | Python | Python |
| Steinbock | CLI Python | R | R | No | No | Python GUI |
| imcRtools | No | No | No | R | R | No |
| SPIAT | No | No | No | No* | R | No |
| Giotto | No | No | R | No | R | No |
| Spectre | No | No | R | R | R | No |
| MCMICRO | nf | nf | nf | No | Python nf | Python GUI |

**Supplementary Table 3:** The availability of essential spatial analysis tools across various computational packages.”

|  | ML cell-type annotation | CN | Spatial Context map | CN interface | Patch proximity | Cell-cell interaction |
| --- | --- | --- | --- | --- | --- | --- |
| **SPACEc** | Yes | Yes | Yes | Yes | Yes | Yes |
| Steinbock | NA | NA | NA | NA | NA | NA |
| imcRtools | Yes | Yes | Yes | Yes | No* | Yes |
| SPIAT | No* | Yes | No | No* | Yes | No |
| Giotto | No | Yes | No* | No* | No | Yes |
| Spectre | Yes | No | No | No | No | No |
| MCMICRO | No | Yes | No | No | No* | No |

## Supplementary Note 1

### Timing

Runtime was estimated using the provided example dataset (2613×2614 px, 58 channels, 50065 cells), except for the tissue extraction steps (28800×23040 px, 29 channels, 147128 cells). The runtime analysis was performed by a researcher who is experienced with the pipeline on a Ubuntu (22.04 CPU: Intel i5 11400 GPU: Nvidia GTX 1070 8 GB RAM: 32 GB DDR4 2400 MHz Storage: 1 TB Gen3 NvMe SSD). Time spent writing new code was not included. The runtime will scale with the complexity and size of the dataset and will depend on the computer used. **Table 1** provides runtime estimates for each step.

### Troubleshooting

**Table 2** provides common errors encountered and their potential reasons and solutions.

**Table 2.**
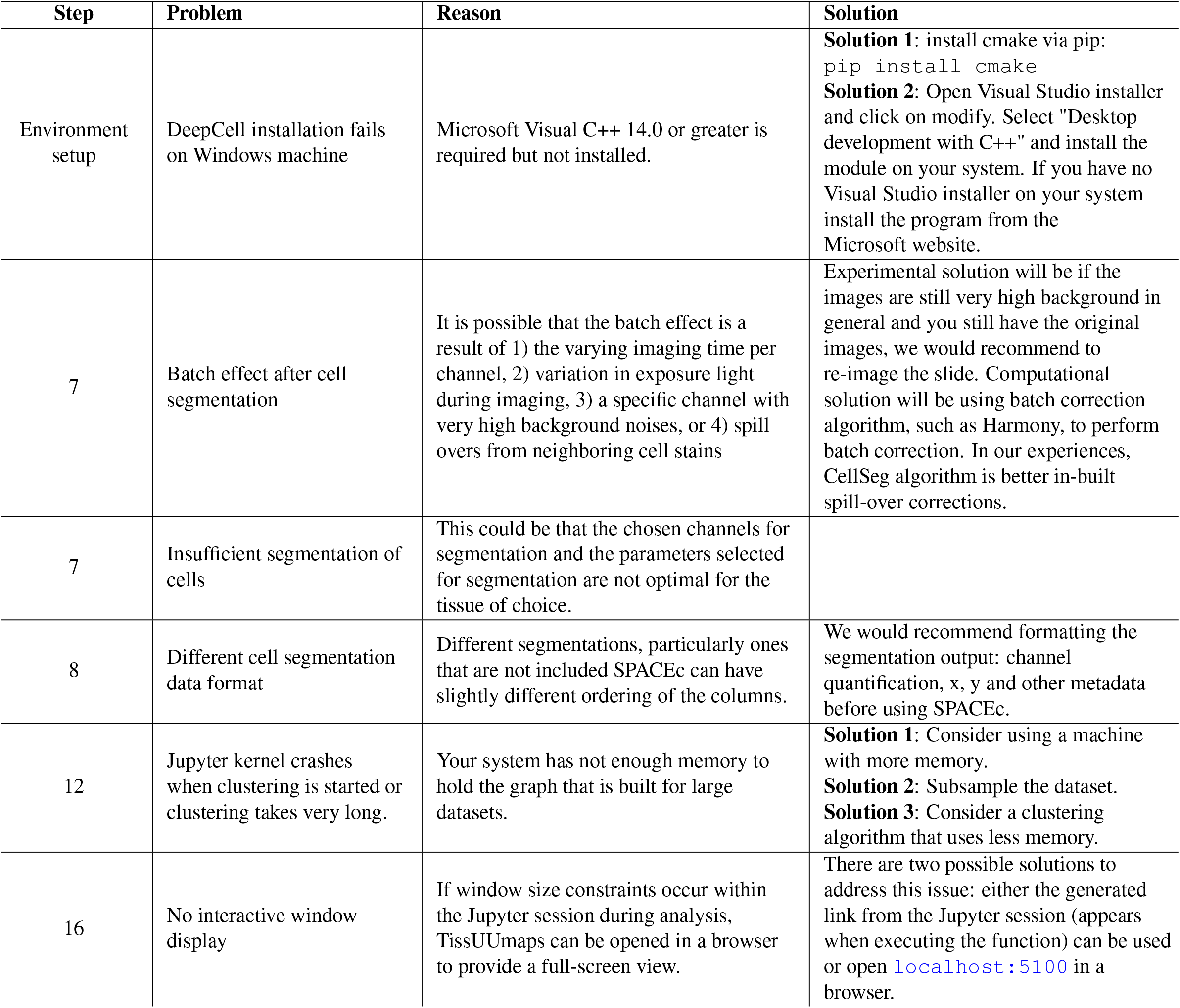
Troubleshooting table.

## Supplementary Note 2

### Software

- SPACEc is compatible with Windows, Linux, and Mac OS (intel/M1/M2).
- SPACEc is implemented in Python version 3.9.0 (https://www.python.org) using Conda version 4.11.0 (https://www.python.org).
- Jupyter Lab version 4.1.1 (https://jupyter.org) is used for data visualization and analysis.

Below, are the versions of the major Python libraries utilized in SPACEc. A yml file is provided on GitHub that lists the software versions for all libraries employed throughout the protocol.

- Anndata 0.10.5
- Cellpose version 3.0.4
- Deepcell version 0.12.9
- HDF5 version 1.14.0
- Keras version 2.8.0
- Scanpy version 1.9.6
- scikit-image version 0.22.0
- scikit-learn version 1.4.1
- TensorFlow version 2.10.0
- Torch version 2.2.0
- TissUUmaps version 3.2.0.5

### Environment setup

1. Install the conda package manager according to instructions available at: https://docs.conda.io/projects/conda/en/latest/user-guide/install/
2. Create and activate a conda environment with Python version 3.9.0

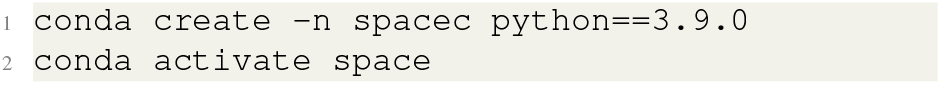
3. Install SPACEc and additional software that is needed to run SPACEc

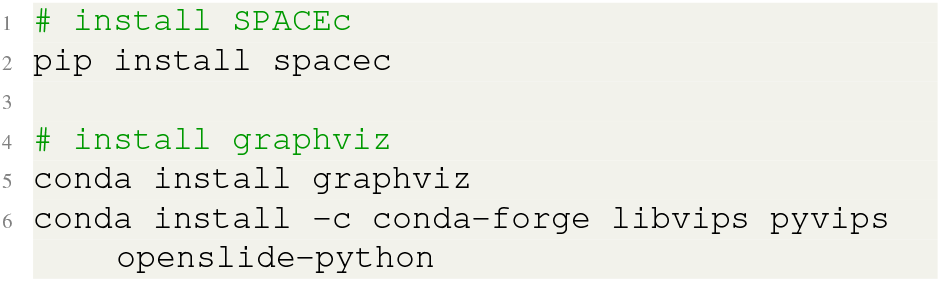
4. After installation, either open a Jupyter Notebook within the environment or activate Python in another integrated development environment

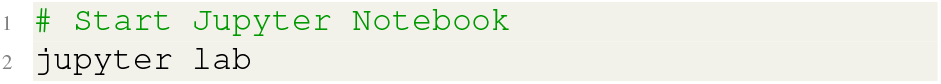

#### A. Tissue image extraction

- **Timing**: 11 min (3.3 GB, 28800x 23040 px, 29 channels, 147128 cells)

The raw output from a PhenoCycler experiment is a qptiff or tiff file. In many cases, multiple tissues are analyzed on the same slide or there are empty spaces around the tissue. For uniform downstream processing as well as optimal use of memory efficiency, first, extract individual tissue images from the qptiff or tiff file and save each as a separate tiff file; these tiff files will be the input for unique coordinates from cell segmentation.

##### A.1. Step 1: Set up the environment

Load all required libraries, set the code, data, and output directory, and define an output path and file name:

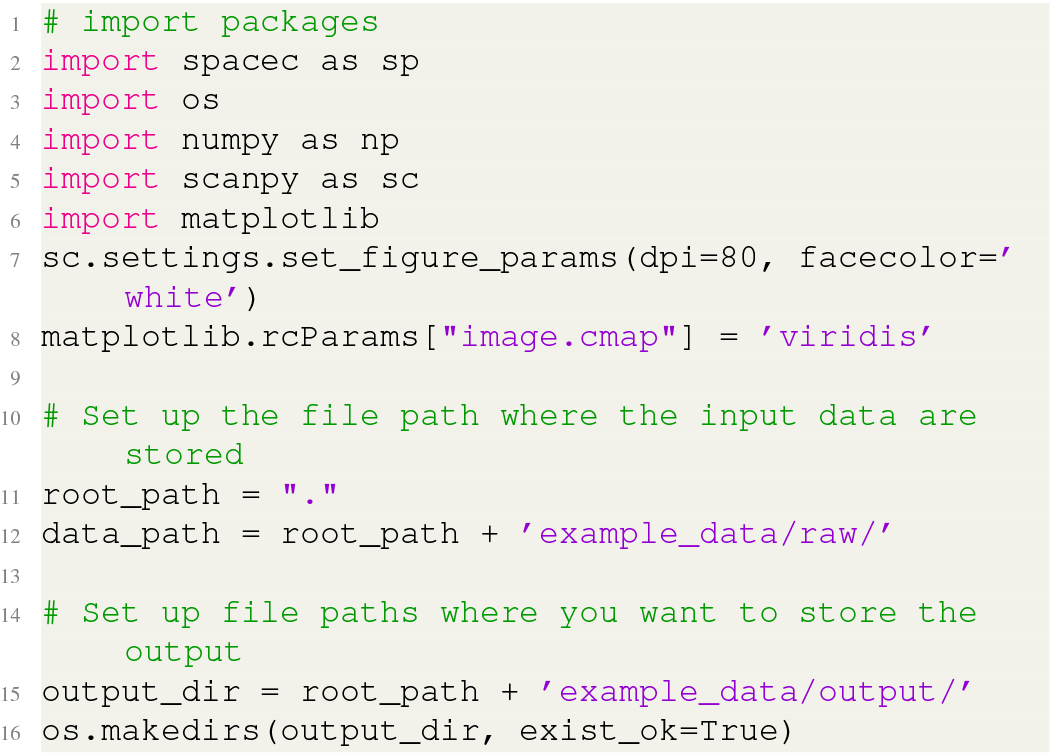

##### A.2. Step 2: Downscale the tissue image

Downscaling of the image is recommended to speed up tissue detection:

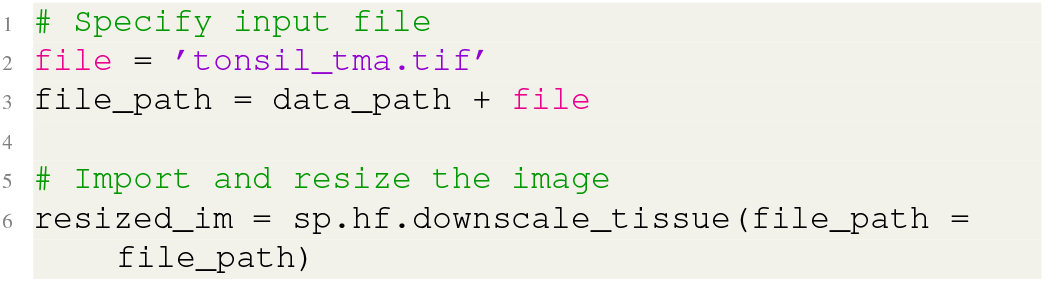

The user should choose cut-offs based on the marker intensity history generated by this function so that the tissue of interest can be appropriately segmented. The cut-off ranges tend to be at the lower quarter of the marker intensity with a typically narrow range of 0.1 between the lower and upper cutoffs.

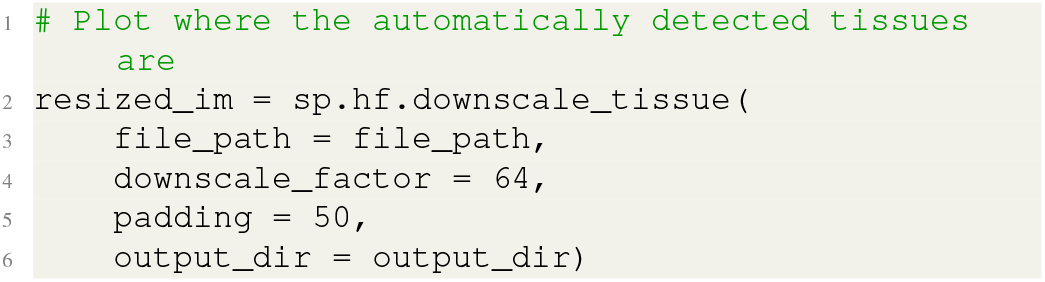

##### A.3. Step 3: Visualize tissue region number

The function below displays the labels of the tissue pieces and the final assignments of regions for the tissue pieces (**Fig. 2A**). Region assignments can be changed through manual input. Users can also combine several multiple disconnected tissue pieces into one coherent piece.

**Supplementary figure 1:**
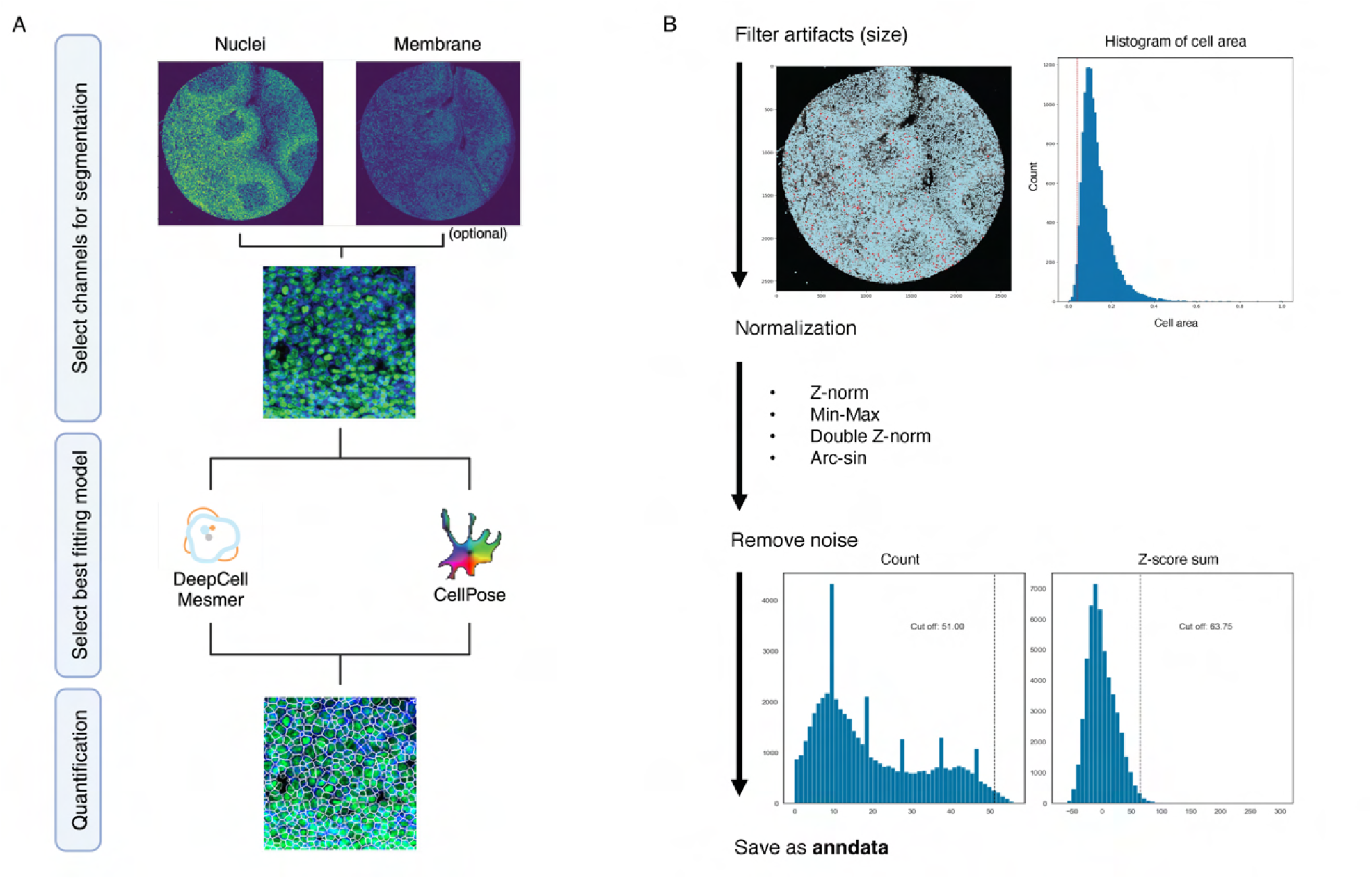
SPACEc preprocessing steps. **(A)** Schematic illustrating how users can choose a combination of nuclei and/or membrane markers to perform segmentation. The user can select one of the three segmentation algorithms included in the pipeline, Cellpose and Mesmer, for segmentation. The segmentation masks can be visualized, and markers quantified. **(B)** Artifacts are removed based on size, guided by the histogram of the size distribution, and staining intensity. After removing the artifacts, there are four different options for data normalization. In a subsequent step, cells that exceed a user-defined threshold for marker positivity are removed. Filtered and normalized data are saved as an Anndata object.

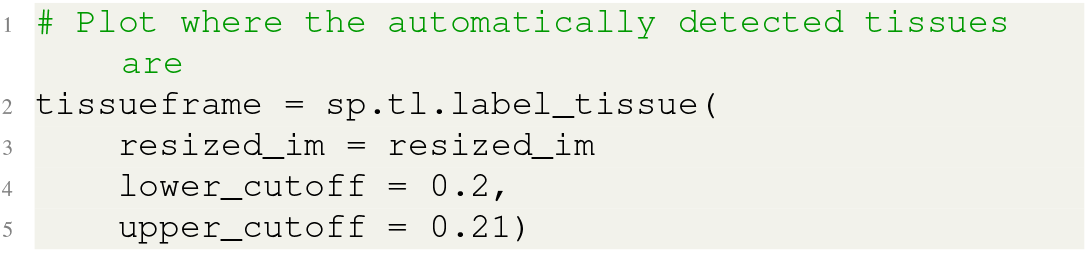

##### A.4. Step 4: Extract individual labeled tissues into separate tiff stacks

The annotated tissue fragments are then cropped from the original input image and stored in the specified output directory as individual tiff stacks. The output file will be formatted as reg001_X01_Y01_Z01.tiff, reg002_X01_Y01_Z01.tiff, etc.

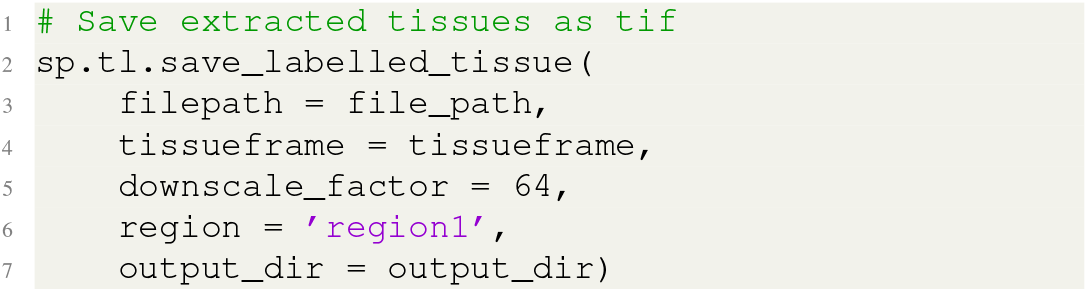

#### B. Cell segmentation and visualization

- **Timing**: 12.8 min (403 MB, 2613×2614 px, 58 channels)

Cell segmentation is performed on extracted tiff files using either Mesmer (33), or Cellpose (34) (**Fig. 2A**). Expression levels for each marker channel are quantified and morphological features such as cell size and shape per cell are calculated.

##### B.1. Step 5: Visualize segmentation channel (optional)

Users may wish to combine multiple channels for membrane or cytoplasm-based segmentation. For instance, in the example data from the tonsil tissue, CD45 and beta-Catenin can be combined to detect cell membranes. The subsequent visualization function enables the user to evaluate the channel selection before initiating the segmentation process.

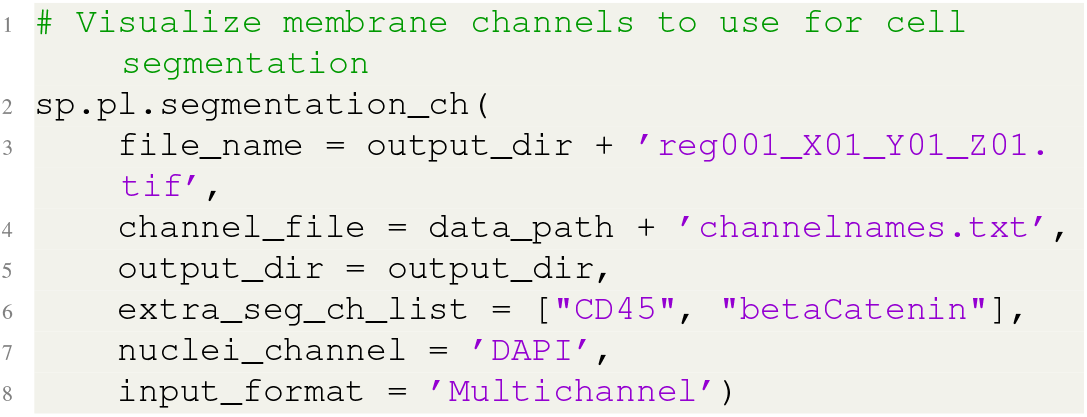

##### B.2. Step 6: Cell segmentation

Next, cell segmentation, marker intensity, and morphological quantification are performed on each cell in the selected image. The function returns the segmentation results in the form of a dictionary that includes both the processed images and the resultant masks. The quantified data are stored as a csv file.

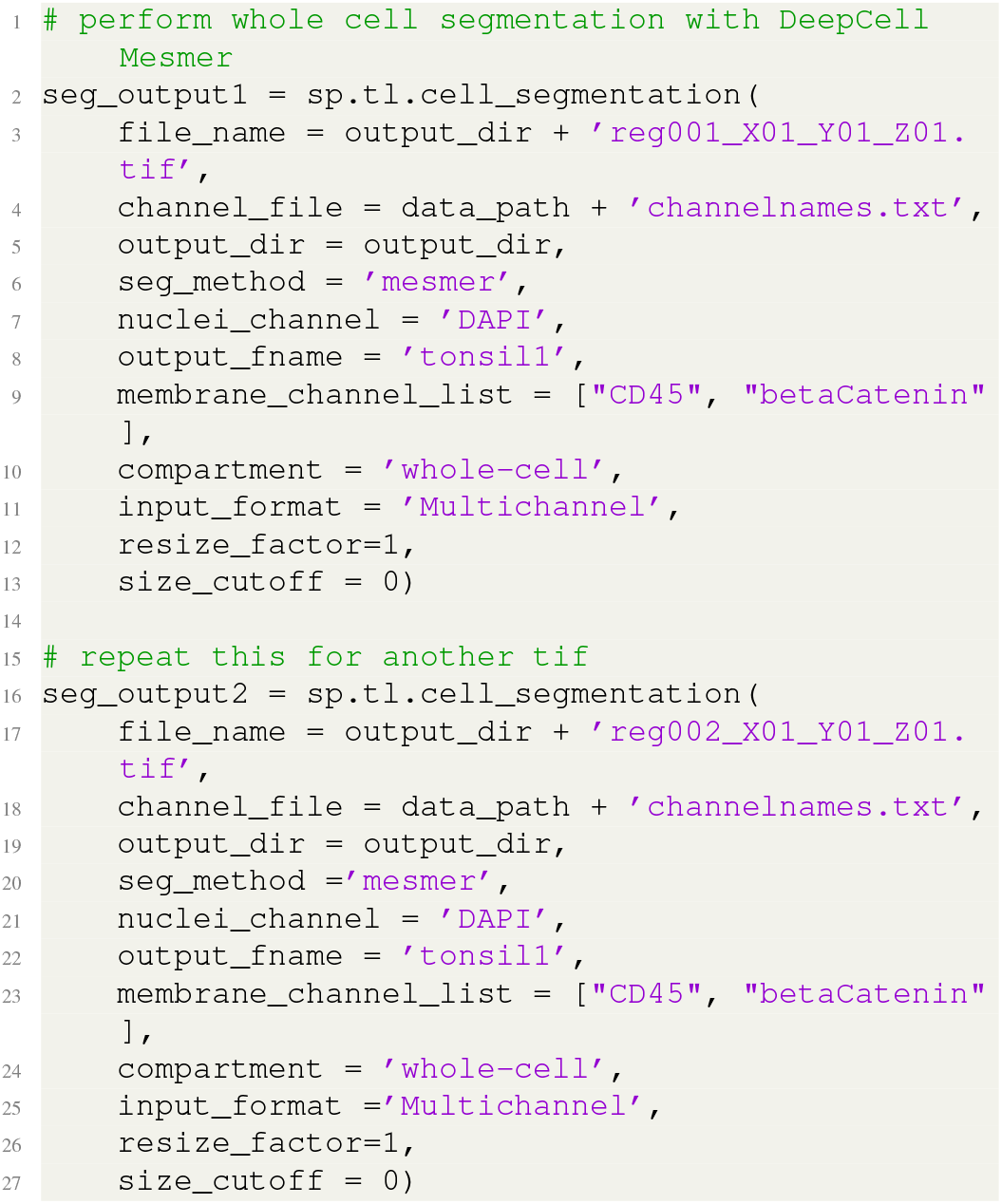

##### B.3. Step 7: Visually inspect segmentation results

Segmentation is crucial for the downstream analysis, so a visualization function is provided that allows the user to inspect the quality of the segmentation results. If the segmentation result is unsatisfactory, the user can repeat the prior segmentation step with different parameters. For example, a different segmentation algorithm or different membrane markers can be used or only nuclei segmentation can be employed. Each segmentation output is saved as a csv file with each row as a single segmented cell with quantified marker expression and x, y coordinates. If the segmentation matches the expected membrane output without over-merging or under-segmenting cells, the segmentation masks are stored as a pickle file.

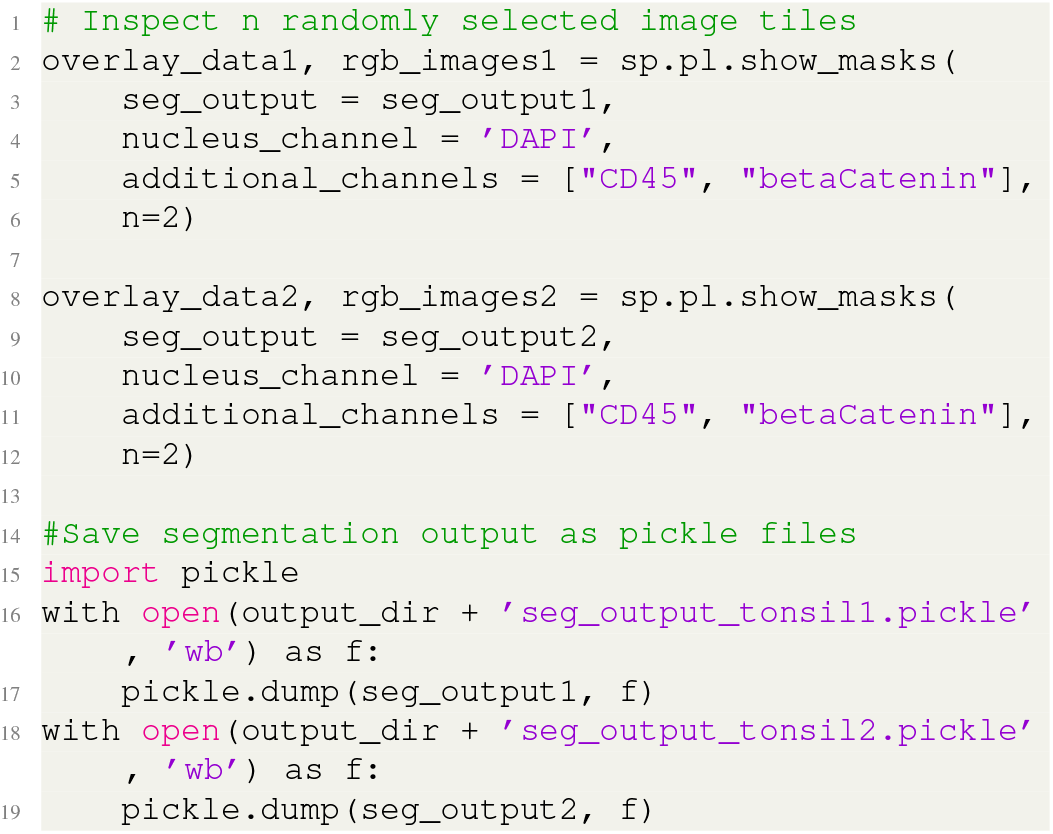

#### C. Data preprocessing and normalization

- **Timing**: 0.3 min (2 csv files (27.5 and 34.5 MB), 52376 cells, 71 features)

After segmenting and quantifying the resultant cell masks, the image data are stored as a single-cell matrix for every segmented image and saved as a csv file. This matrix saves cells as rows and features (marker intensity values and metadata) as columns. Data preprocessing and normalization steps allow the user to combine the segmentation results into a single file, perform filtering and normalization, and generate an Anndata object that contains the single-cell information in a standardized format.

##### C.1. Step 8: Load the segmented results

Segmented csv files are loaded and concatenated as one data frame for further processing. The example uses two tonsil images. Meta information, such as region name (in string format, e.g. “reg001”, “reg002”) and image information (in string format, e.g. “tonsillitis”, “tonsil”), is required to discern individual segmented images.

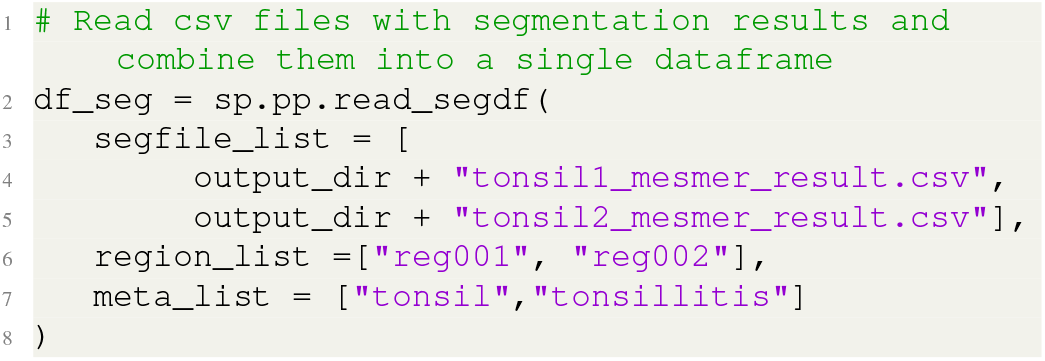

##### C.2. Step 9: Initial removal of artifacts and debris

DAPI intensity and cell size are used to identify and remove artifacts such as dust particles and cell debris. Based on our experience with CODEX analysis, only about 1 % of cells should be removed based on size and nuclear intensity thresholds. Users can also visualize the cells that have been removed to determine whether the filtering step was effective (see *Step 16*).

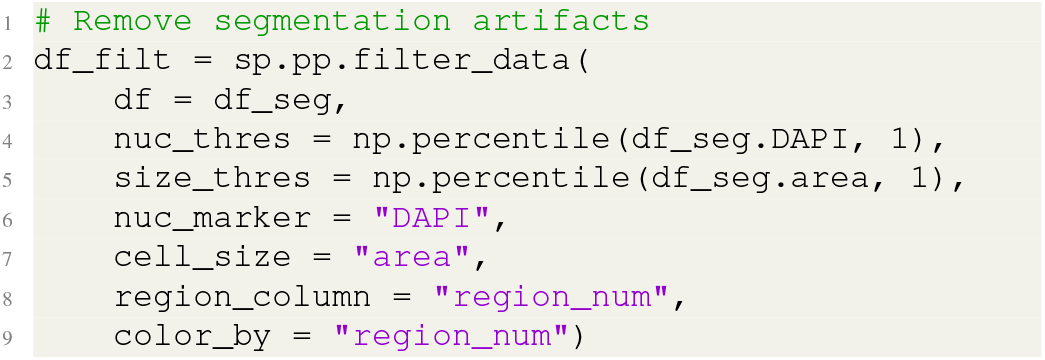

##### C.3. Step 10: Normalize data

Normalizing marker intensity reduces the intensity ranges between channels and provides more accurate cell-type annotation. We have previously shown that Z-score normalization is the most robust(35). To ensure that only marker expression is normalized, metadata column names of information that should not be included in the normalization are placed in list_out and those to be included are placed in list_keep. SPACEc provides “double_zscore”, “MinMax”, and “ArcSin” normalization. The detailed documentation of each normalization can be found in the *Supplement Notes*.

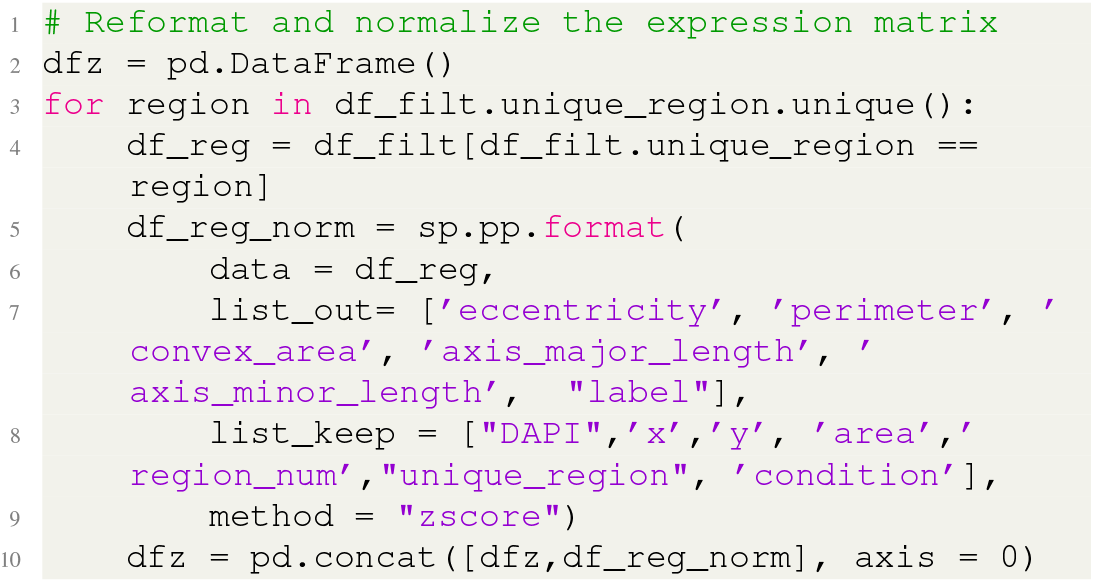

##### C.4. Step 11: Second round of filtering for aberrantly stained cells

A second round of filtering is used to remove cells that stain positive for many markers. The first function generates two histograms that allow the user to adjust the threshold and inspect the data. A suitable threshold is selected and then applied to the data. As an example, in our experience if 50 markers are used, cells with summed z scores of above 35 that have z scores of more than 1 for 35 or more markers should be removed. This generally does not eliminate more than 1 percent of the data. It is acceptable to be conservative at this step because unsupervised clustering should also remove these aberrant cells. If certain regions of an image have many flagged cells, it can be helpful to visualize the “removed cells” by plotting in spatial coordinates with catplot function using the returned “cc” dataframe from the sc.pp.remove_noise pipeline. The denoised data are then saved as an Anndata object. Through the inspection of marker intensities, users can exclude the markers that did not work or did not stain as expected by adding them to nonFuncAb_list.

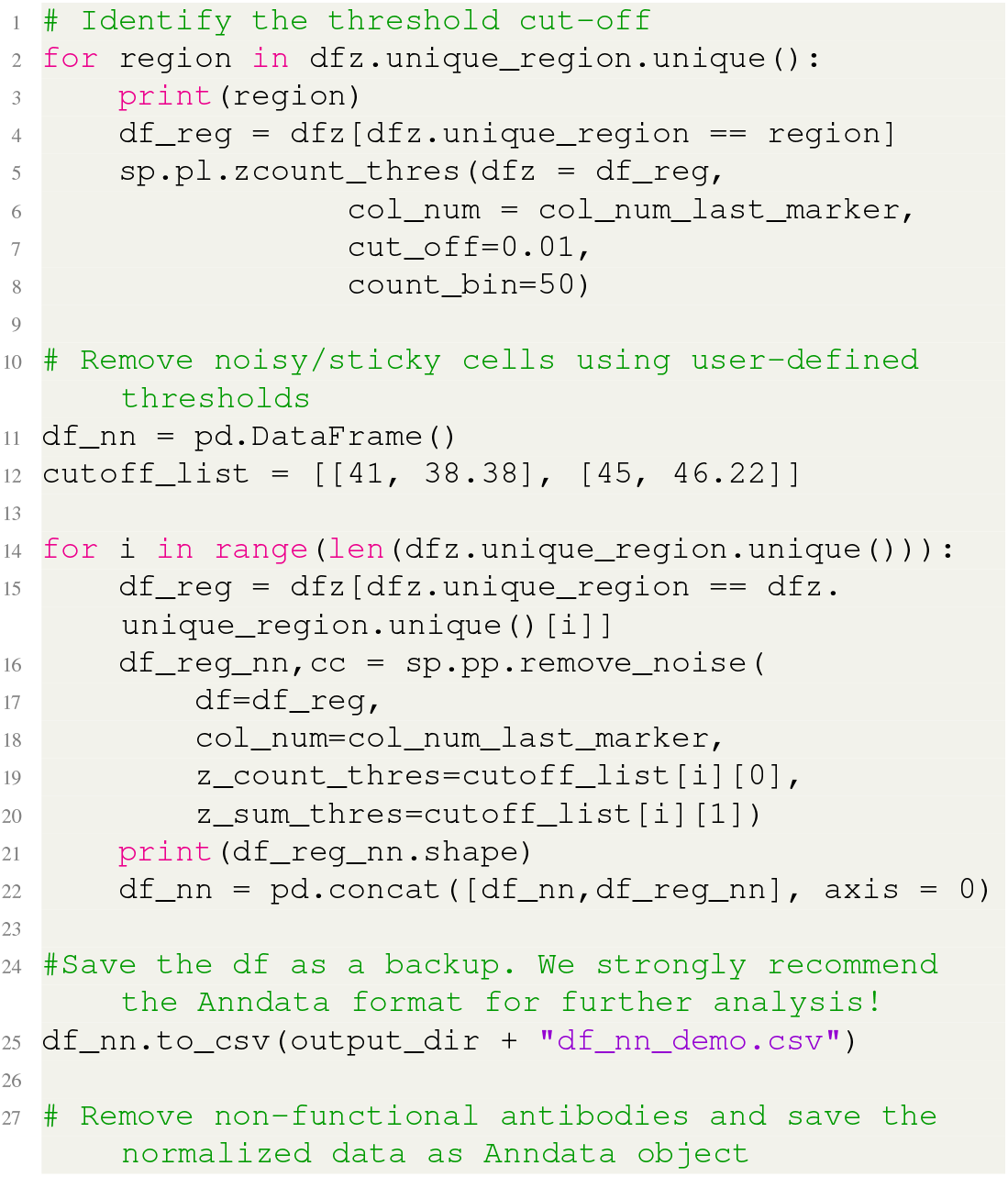
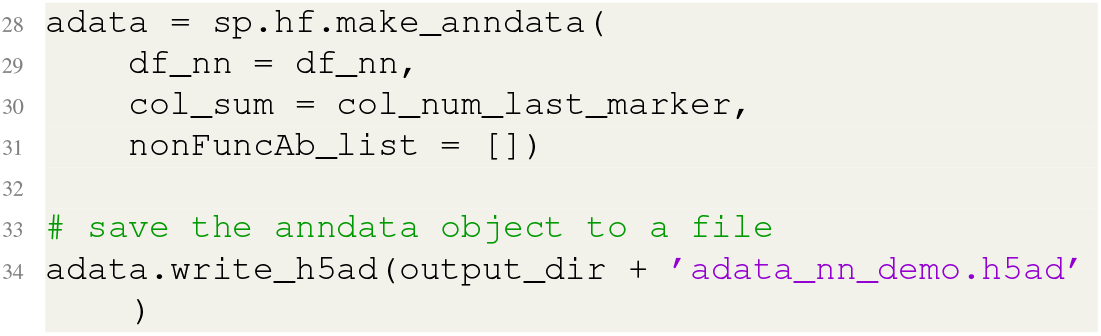

#### D. Cell-type annotation

Option A: Unsupervised clustering

- **Timing**: 2 min (52376 cells, 39 features) plus time for manual annotation of the clusters)

After preprocessing the single-cell data, the next step is to assign cell types. One of the most common approaches to identifying cell types is through clustering. SPACEc uses Louvain and Leiden clustering (45). For complex datasets, multiple rounds of iterative clustering are often necessary (for example, see our GitHub repository). There are two typical approaches to clustering of multiplexed imaging: 1) coarse clustering at the broad cell-type level (e.g., B cells, T cells) followed by subclustering or 2) overclustering and then merging similar clusters. In this protocol, we describe the first approach, although overclustering can also be implemented from within SPACEc.

##### D.1. Step 12: Cell-type annotation via clustering

First, unsupervised clustering is used to group cells, and then specific marker expression is used to annotate each cluster. All markers or a subset of markers can be selected for clustering. In our experience, clustering on cell-type specific markers (e.g., CD3) and excluding functional markers (e.g., PD-1) in the initial clustering yields better results.

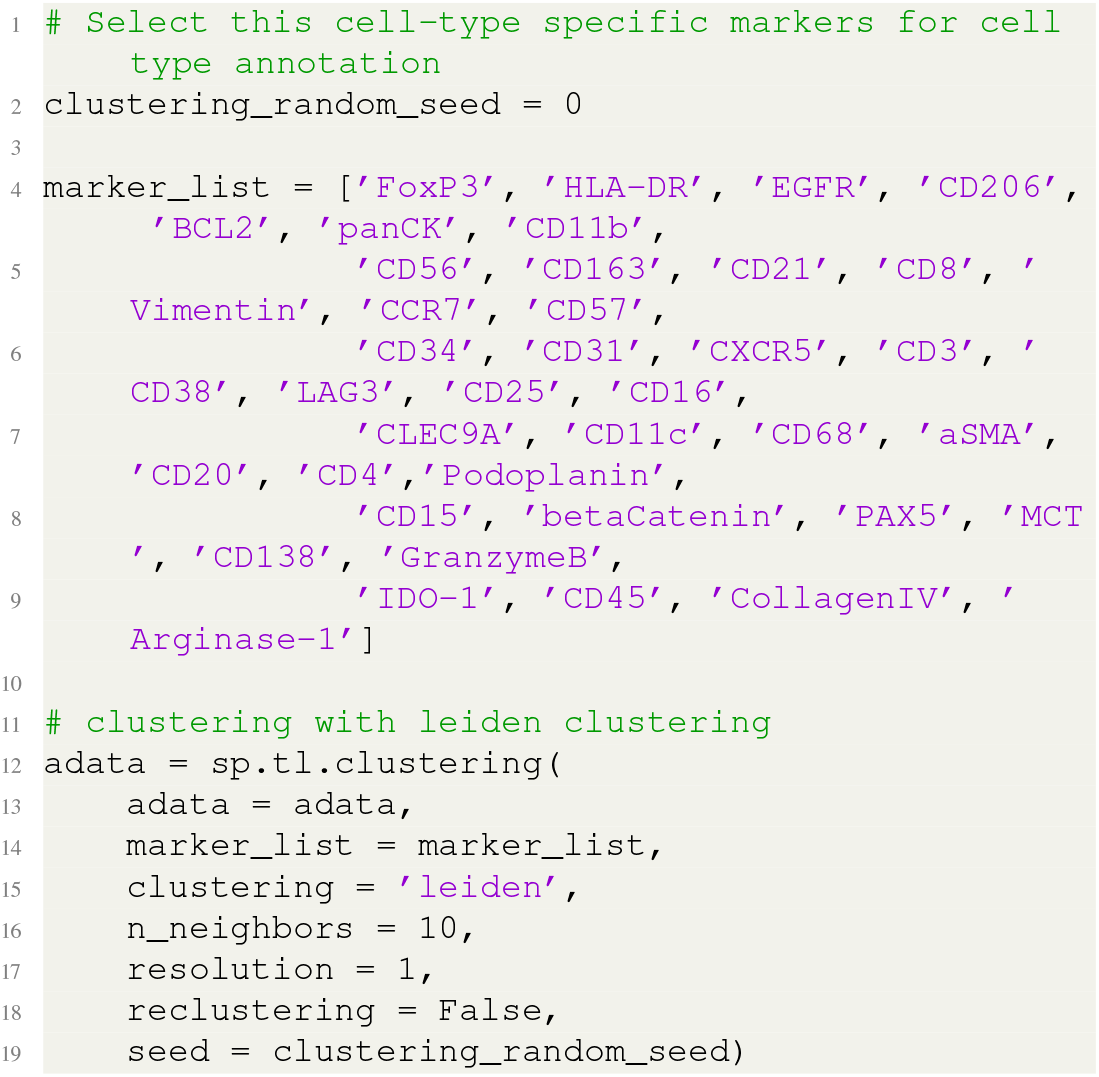

##### D.2. Step 13: Visualize and annotate clustering results

The visualization output of the clustering result for the example data is shown in **Supp. Fig. 2A-B**. The UMAP visualization facilitates the assessment of the separations between clusters and their relationships, and the dotplot reveals the enrichment of specific markers for each cluster.

**Supplementary figure 2:**
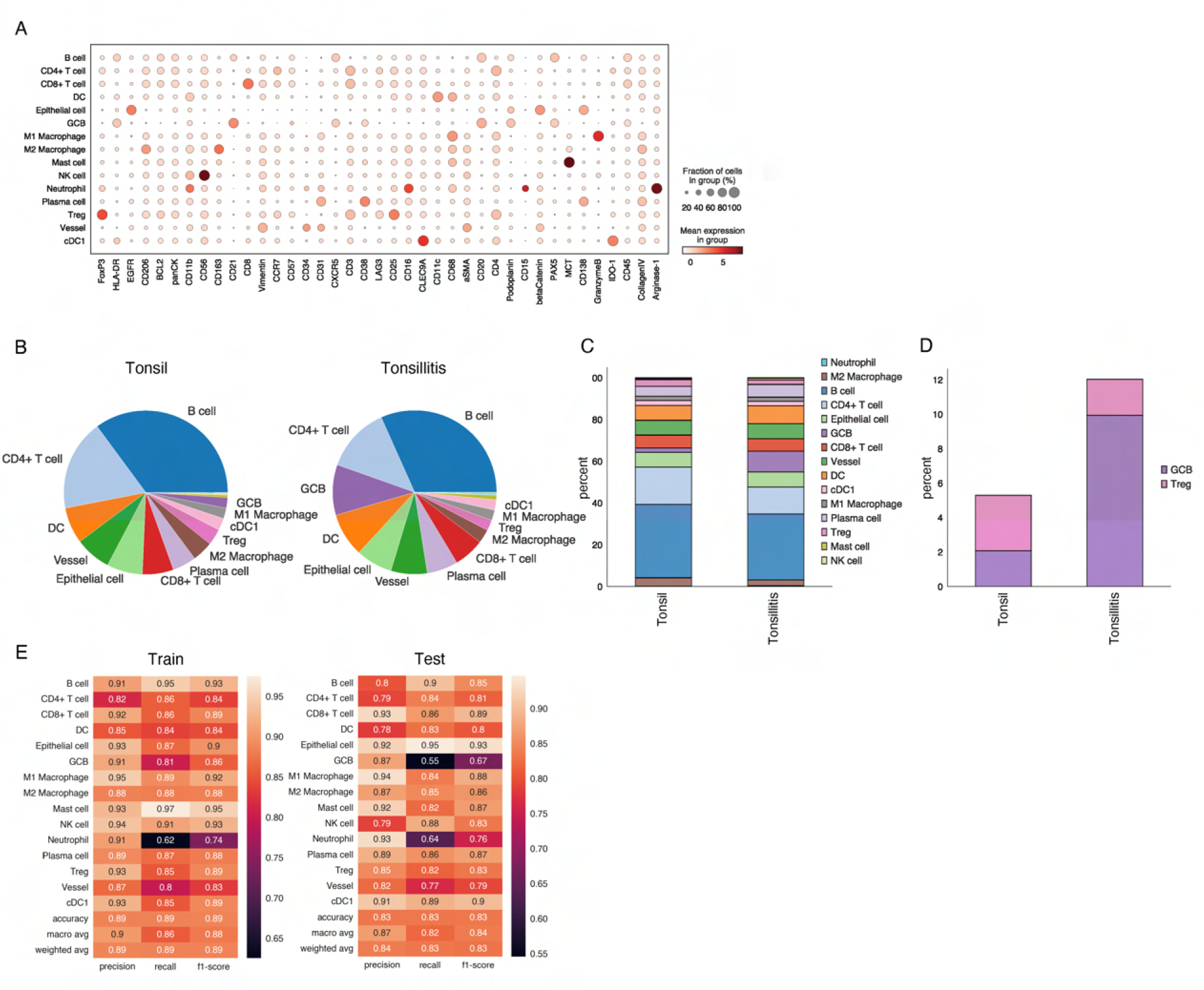
Cell-type annotation and analysis. **(A)** A dotplot shows the marker expression (x-axis) for each cell type (y axis). **(B)** Cell-type percentages for healthy tonsil (left) and inflamed tonsil (right) visualized as a pie chart. **(C)**SPACEc also enables visualization of cell-type percentages as a stacked bar chart. **(D)** Percentages of users’ specific cell types can be displayed in a stacked bar chart. **(E)**Tables on the performance of the SVM model on training and test data. Each row represents a cell type. The columns are evaluation metrics of precision, recall, and f1-score, which are measurements of the classification performance and ranges from 0 to 1. The higher the numeric number, the better the SVM performance.

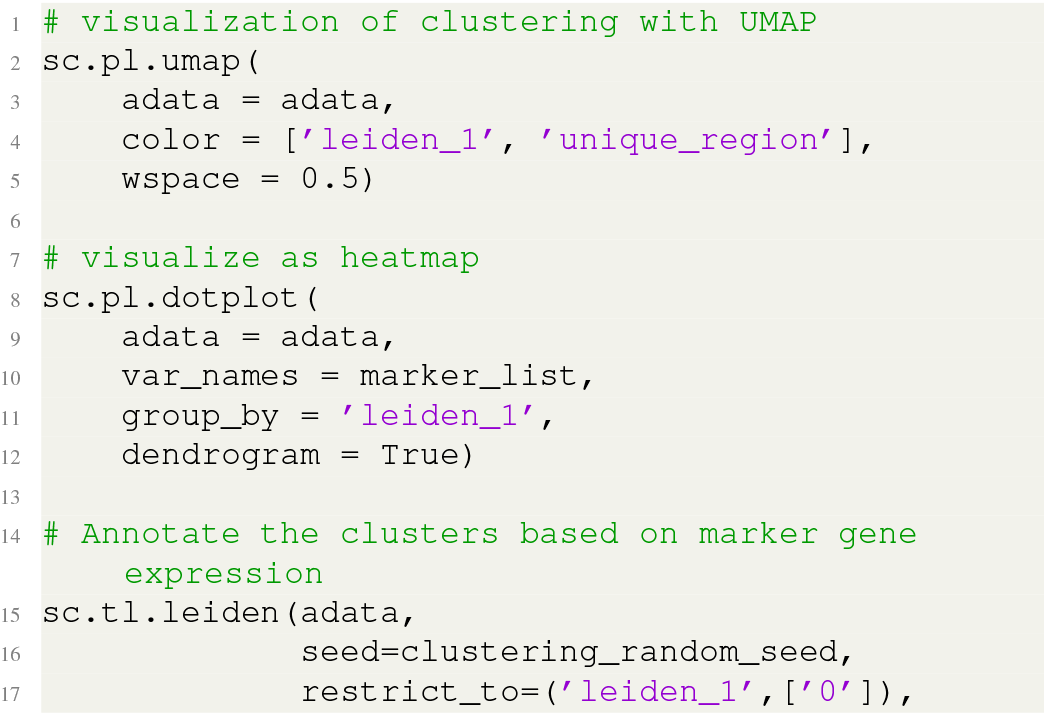
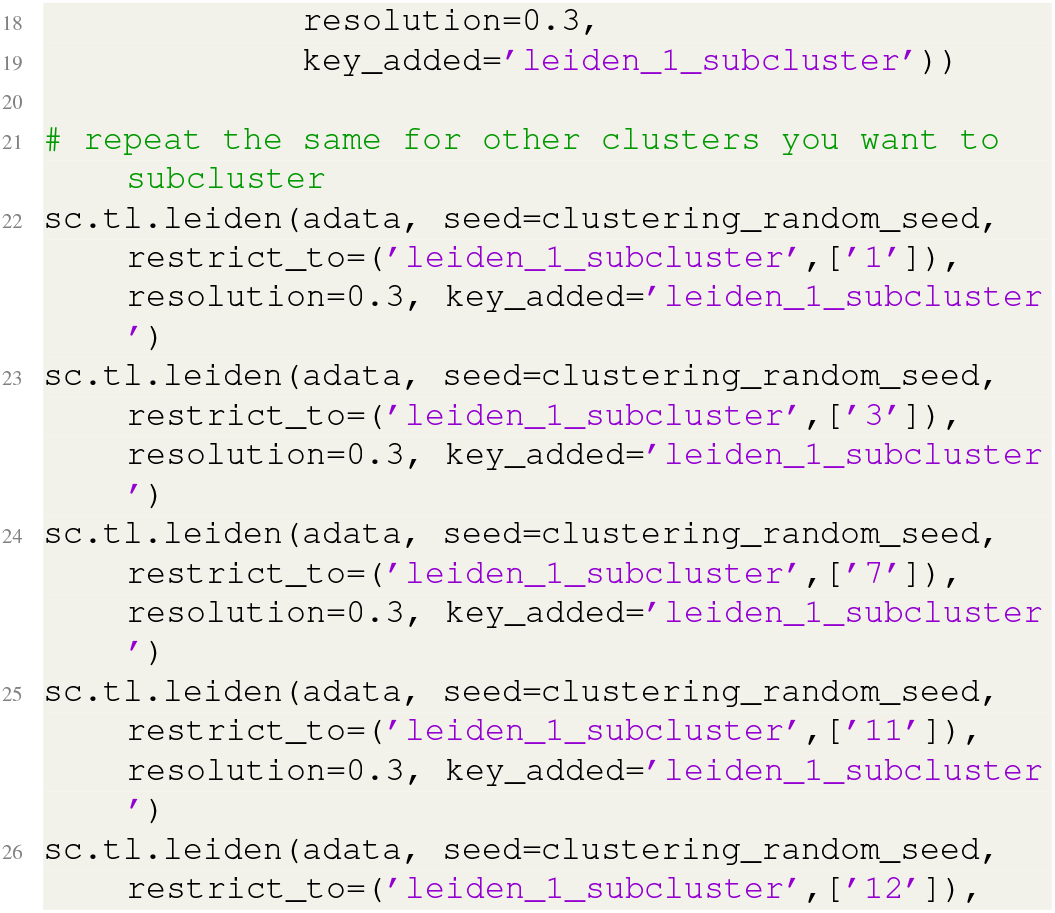
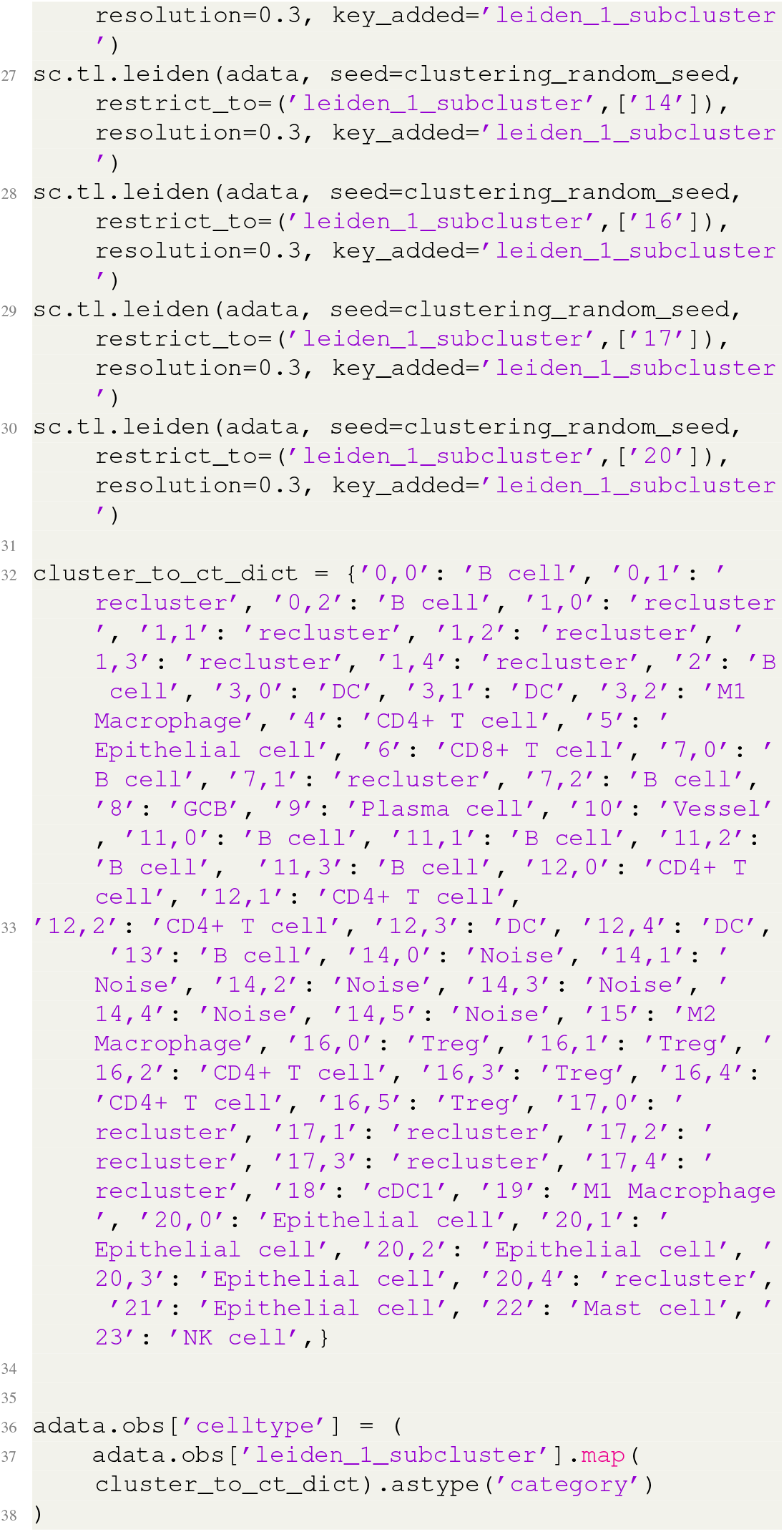

Visualizing the annotated cell types in their original tissue coordinates is crucial to the determination of the identities of each cluster. SPACEc provides a collection of functions for various visualization types (**Supp. Fig. 2C-E**). The annotation results can also be explored as interactive visualization with TissUUmaps. Visualization validates the results of clustering this allows annotations to be compared to molecular marker expression.

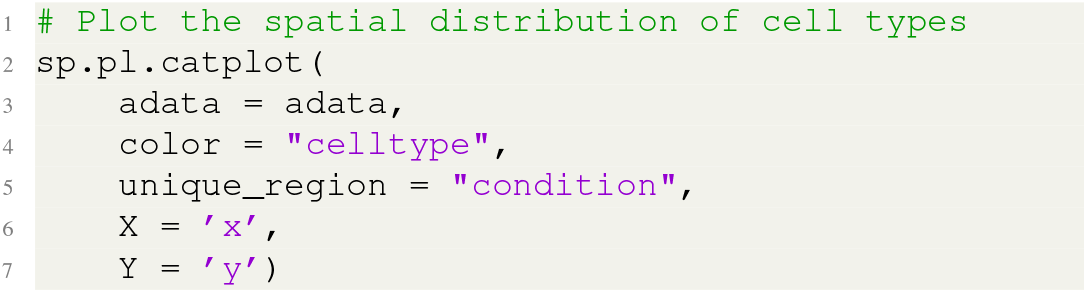

##### D.3. Step 14: Compute basic statistics of cell-type composition

SPACEc can provide and plot basic statistics on the cell-type abundances (**Supp. Fig. 2D-E**)

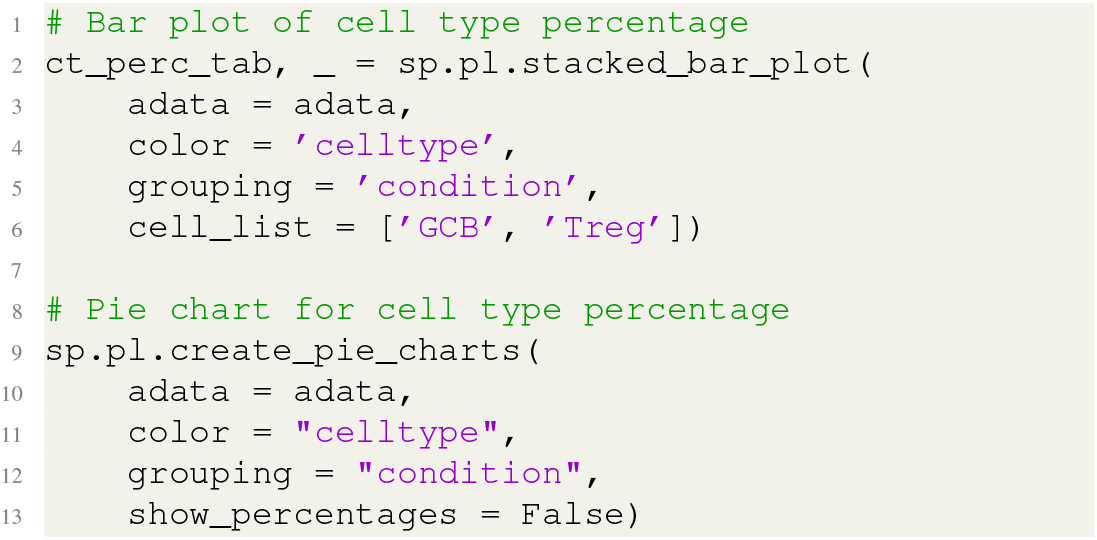

Option B: Supervised machine learning-based cell-type annotation

- **Timing**: 13.6 s (52376 cells, 39 features)

Clustering can be very time-consuming, and if a dataset with similar (preferably the same) marker panels and imaging conditions has been previously annotated, machine learning approaches can be a time-efficient way to transfer labels to an unannotated dataset. SPACEc incorporates an SVM-based approach for machine learning-guided cell-type annotation. The method allows cell-type assignment in unannotated samples based on reference data.

##### D.4. Step 15: Cell-type annotation via machine-learning classification

To train an SVM, one needs previously annotated data as a training dataset. For demonstration purposes, we use annotated tonsil data as training and the unannotated tonsillitis data as validation (**Supp. Fig. 3F**). First, the annotated and training data are loaded

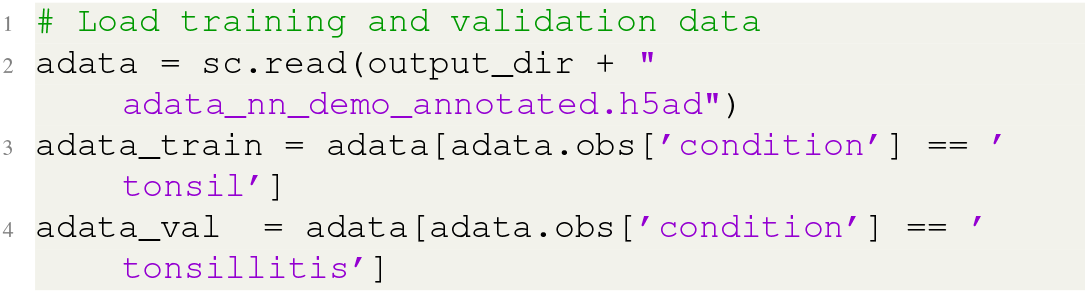

Second, the SVM model is trained on the previously annotated multiplexed imaging data:

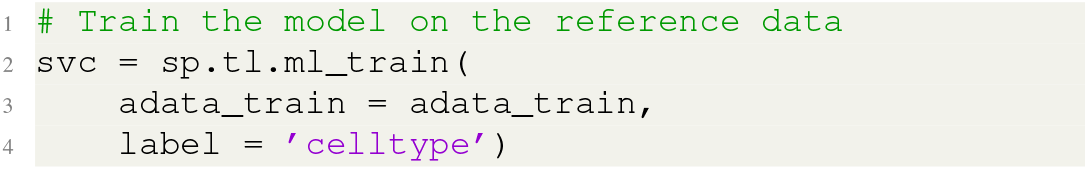

Third, the SVM model is used to predict cell-type labels for the validation dataset:

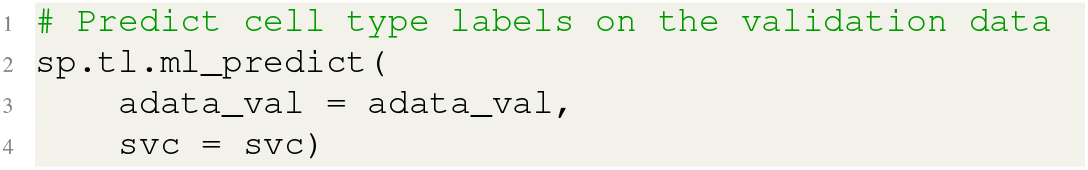

Finally, the results are inspected. Since we know the ground truth for the tonsillitis data, we were able to evaluate the accuracy of the machine learning-based approach.

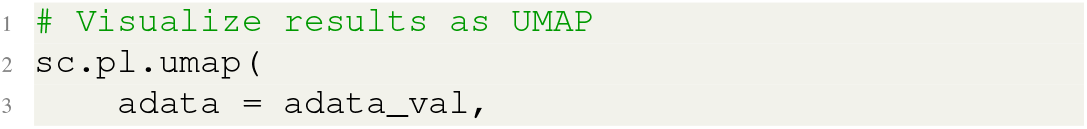
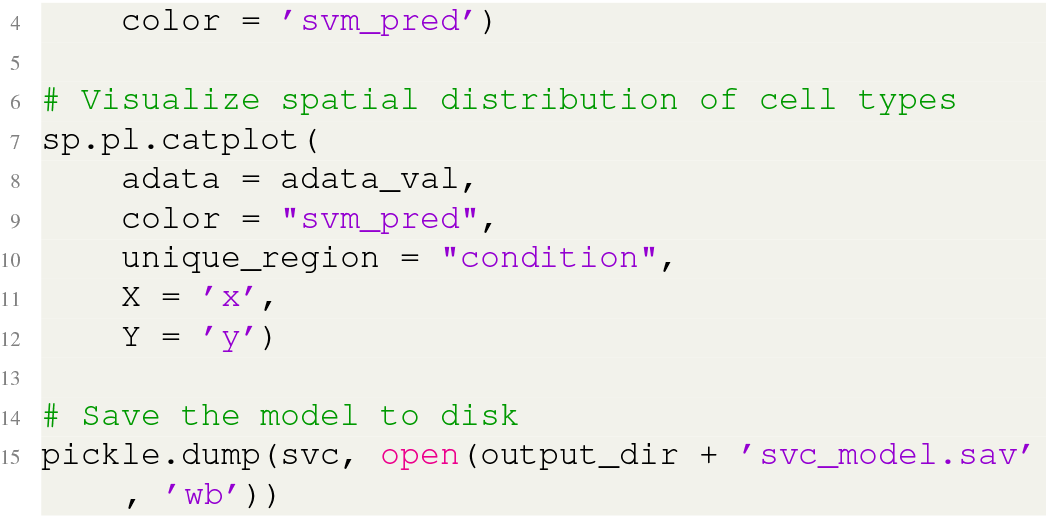

#### E. Interactive data inspection and exploration

- **Timing**: 2.9 s (annata with 52376 cells with 57 features, a tiff image of 403 MB, 2613×2614 px)

Users can inspect the quality of the annotation interactively using TissUUmaps (**Fig. 2D** and **Supp. Fig. 3**), which enables interactive visualization of segmentation results, spatial cell distribution, and delineation of regions of interest.

##### E.1. Step 16: Prepare data for an interactive session

To utilize the interaction session from SPACEc, the Anndata data (generated in *step 11*), the original multiplexed images, and stored segmentation results (generated in *step 6*) are used as input. Additionally, the user needs to indicate a column containing the region information within the Anndata object (region_column) and a selected region. Note that the indicated region must match the images saved in the pickle file. Lastly, the user provides the names of the columns that contain the spatial coordinates as well as a column by which to color the cells (usually cells are colored by cell type).

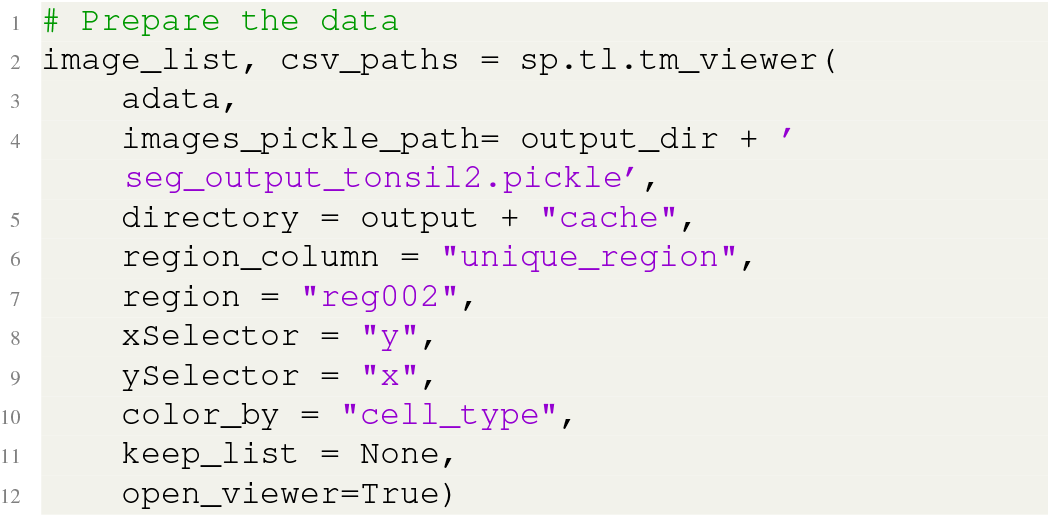

#### E.2. Step 17: Additional interactive data exploration (optional)

TissUUmaps offers various data exploration options such as annotation of regions, analysis of the cellular composition, highlighting regions with similar compositions, and visualizing the relationships between feature space and the image. Here we showcase a few standard plug-ins, but additional plug-ins are available on the TissUUmaps official website (40). To visualize the cellular composition in a specific area, the user can directly draw and mark a region on the image and then apply the Plot_Histogram plug-in (**Supp. Fig. 3C**). The Points2Regions plug-in can be utilized to identify and highlight regions of similar composition (**Supp. Fig. 3B**). The Feature_Space plug-in allows the user to directly draw on the dimension-reduced feature space and to pinpoint cells within the image based on their position in the feature space (**Fig. 2D**). The Spot_Inspector plug-in visualizes a cell, cell type, or region and inspects the marker expression within the original multiplexed images (**Supp. Fig. 3D**). This can help validate the cell-type annotation.

##### F. Spatial analysis

- **Timing**: 1.4 min (anndata with 52376 cells with 57 features)

Cellular neighborhood analysis involves four main steps (**Fig. 2C**): First, a window is drawn around each cell, encompassing the centered cell and its n nearest neighbors. For instance, if a window size of 10 is chosen, each window contains the central cell and its nine nearest neighbors (**Supp. Fig. 4A**). In the second step, the cell-type composition per window is computed and stored as a vector. Third, the cell-type composition vectors per window are clustered into a user-defined number. Finally, the identity of each cellular neighborhood (CN) is assigned based on relative cell-type enrichment.

##### F.1. Step 18: Compute cellular neighborhoods

The x, y co-ordinates and cell types are used to compute CNs. Users should experiment with different combinations of window size and number of neighboring cells particularly when the analysis is of a new type of tissue or addresses a novel biological question. Smaller window sizes (k) will result in the identification of smaller-scale structures that likely represent dense areas of cells with similar cell types. In general, we start with k values of 5, 10, and 20 and evaluate how CN assignments differ across these ranges. A range of numbers of neighboring cells should also be evaluated to identify the value where overclustering is detected. Generally, in a dataset with 20 to 40 cell types, approximately 20 CNs are reasonable. However, the window size and number of neighborhoods depend on the depth of the data and the biological question of interest. The function below outputs an elbowplot plot based on the distortion score, which shows how well the CNs are clustered to provide guidance on parameter selection.

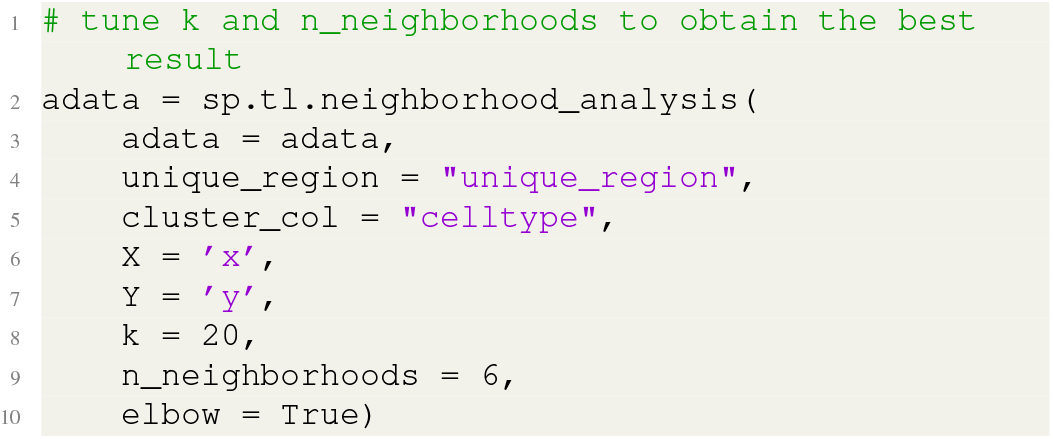

##### F.2. Step 19: Visualize and annotate the cellular neighborhoods

To analyze and annotate CNs, we visualized the cell-type enrichment and the spatial distribution of the CNs (**Supp. Fig. 4B-C**). We first generate a unique and reusable color dictionary to create a consistent color representation of the CNs. A random color palette will be generated if a user does not specify a dictionary. Next, a heatmap is generated to evaluate the enrichment of cell types per CN. This heatmap can help identify the multicellular structures represented by a CN since the heatmap reveals both depletion and enrichment of cell types. For example, a vasculature CN may include endothelial and smooth muscle cells but would exclude epithelial cells. Plotting on the x, y coordinates reveals whether the localization of the CNs corresponds to known tissue units.

**Supplementary figure 3:**
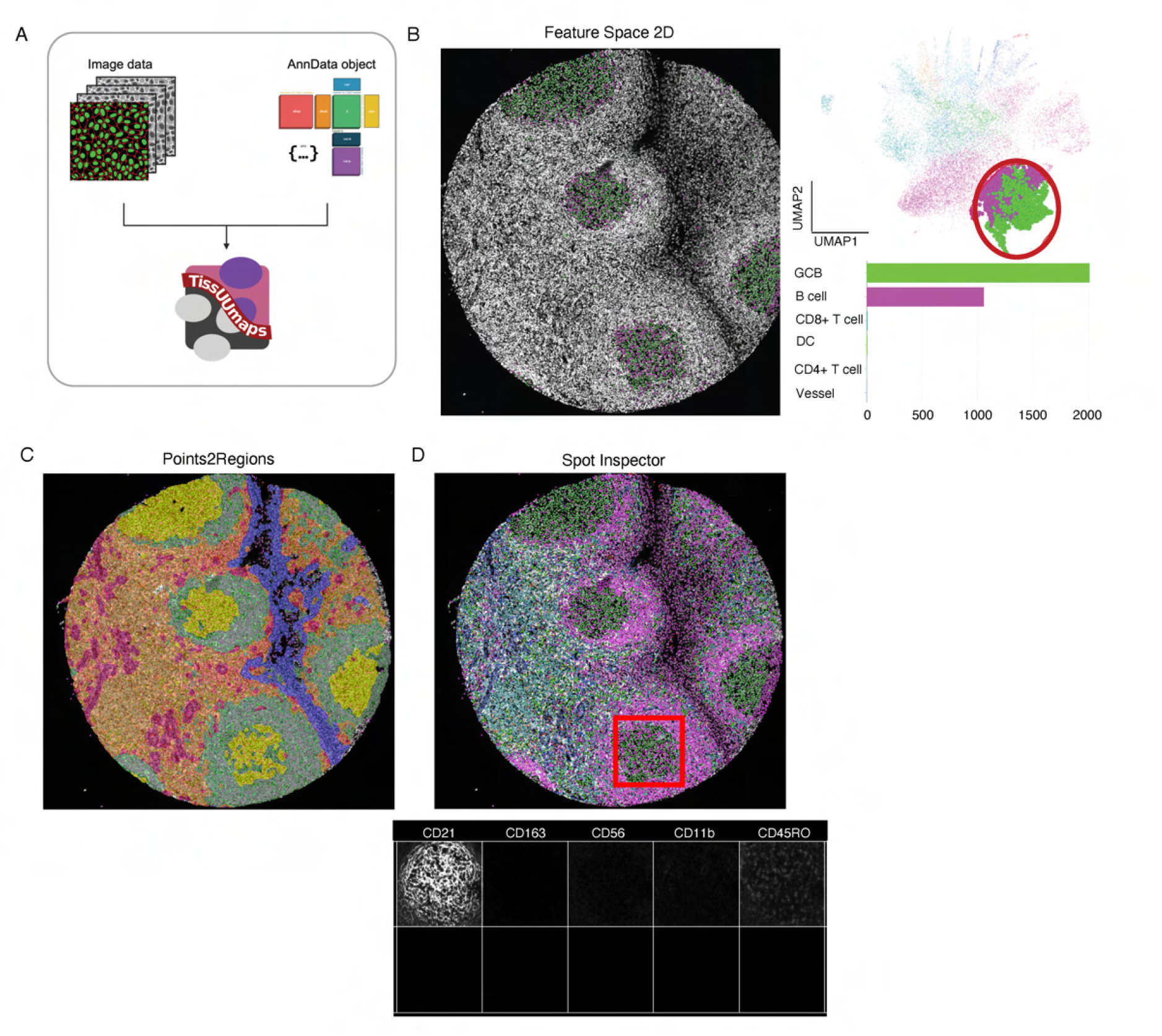
Interactive data visualization and exploration. **(A)** SPACEc imports raw multiplexed images and Anndata objects and uses TissUUmaps to overlay the metadata stored in the Anndata object on the raw multiplexed images. **(B)** The Feature Space 2D plug-in enables users can select a group of cells (e.g. the green and purple cluster) from the pre-computed UMAP space or other pre-computed dimensionally reduced feature space (right). The selected cells are located on the raw multiplexed images will be shown (left) as well as the cell type composition within the circled area (bottom left). **(C)** The Points2Regions plug-in allows users to identify regions/cellular neighborhoods of similar composition after specifying regions/cellular neighborhood numbers. **(D)** The Spot Inspector plug-in can be used to visually inspect marker expression in selected areas.

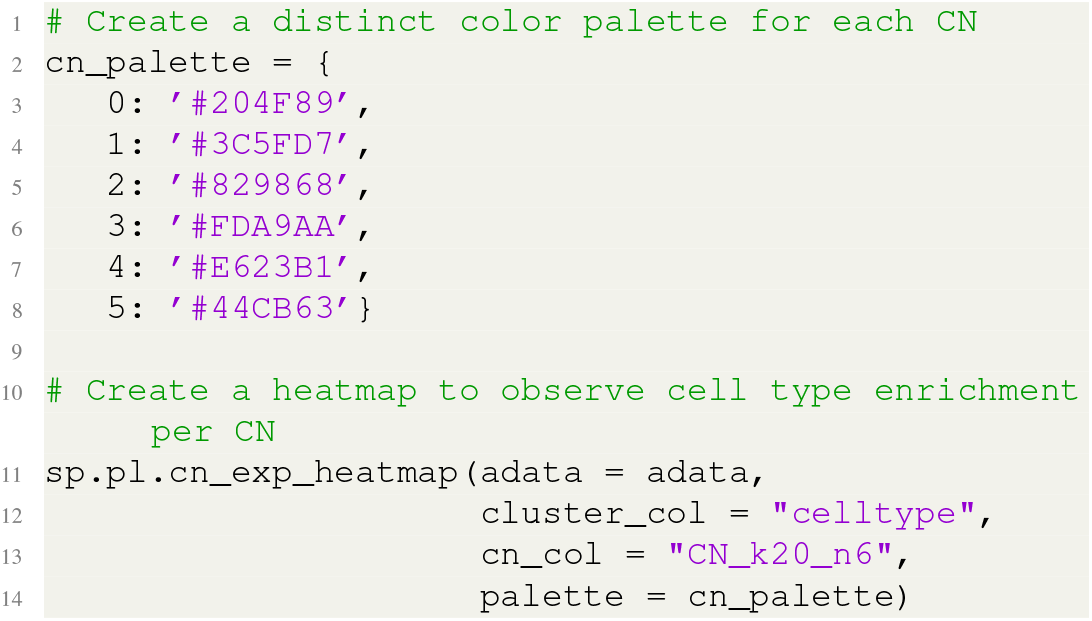

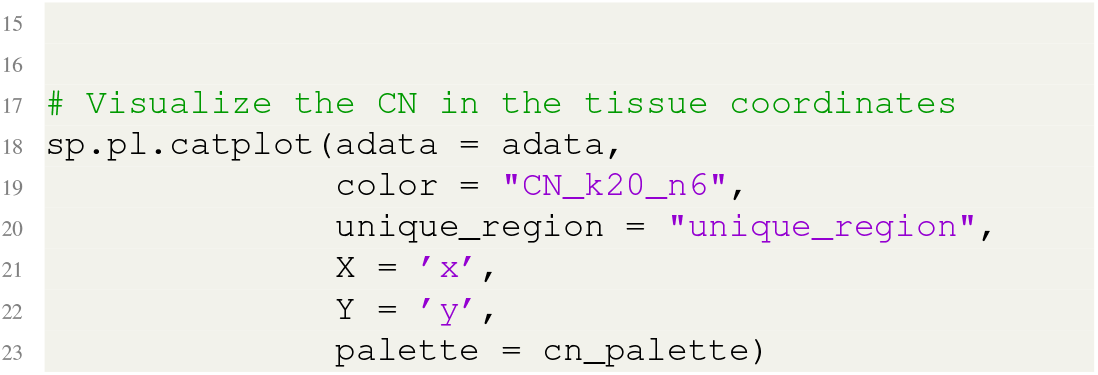

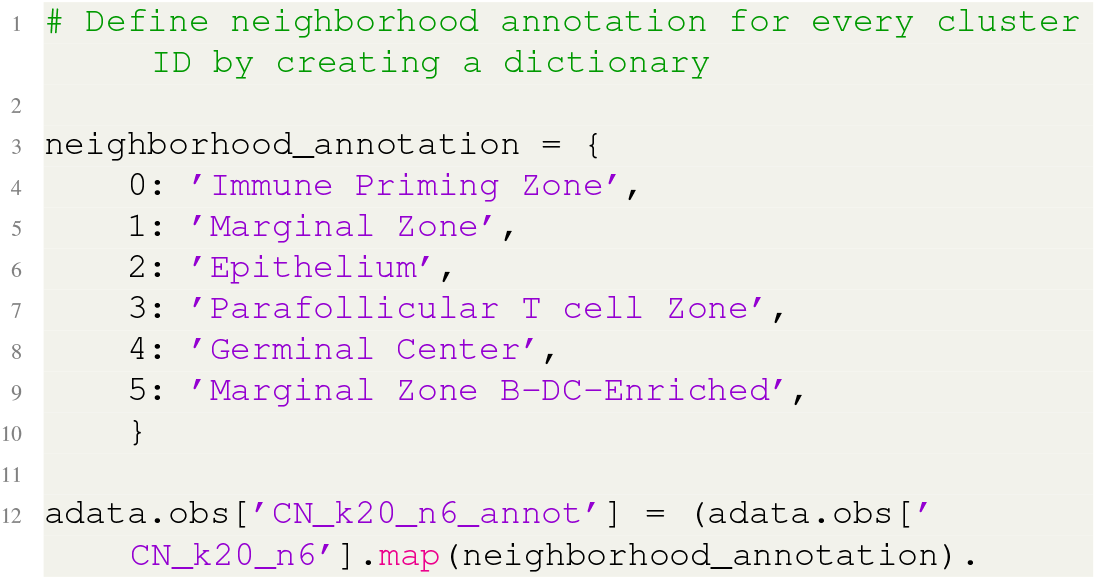
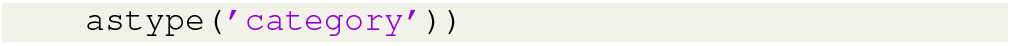

**Supplementary figure 4:**
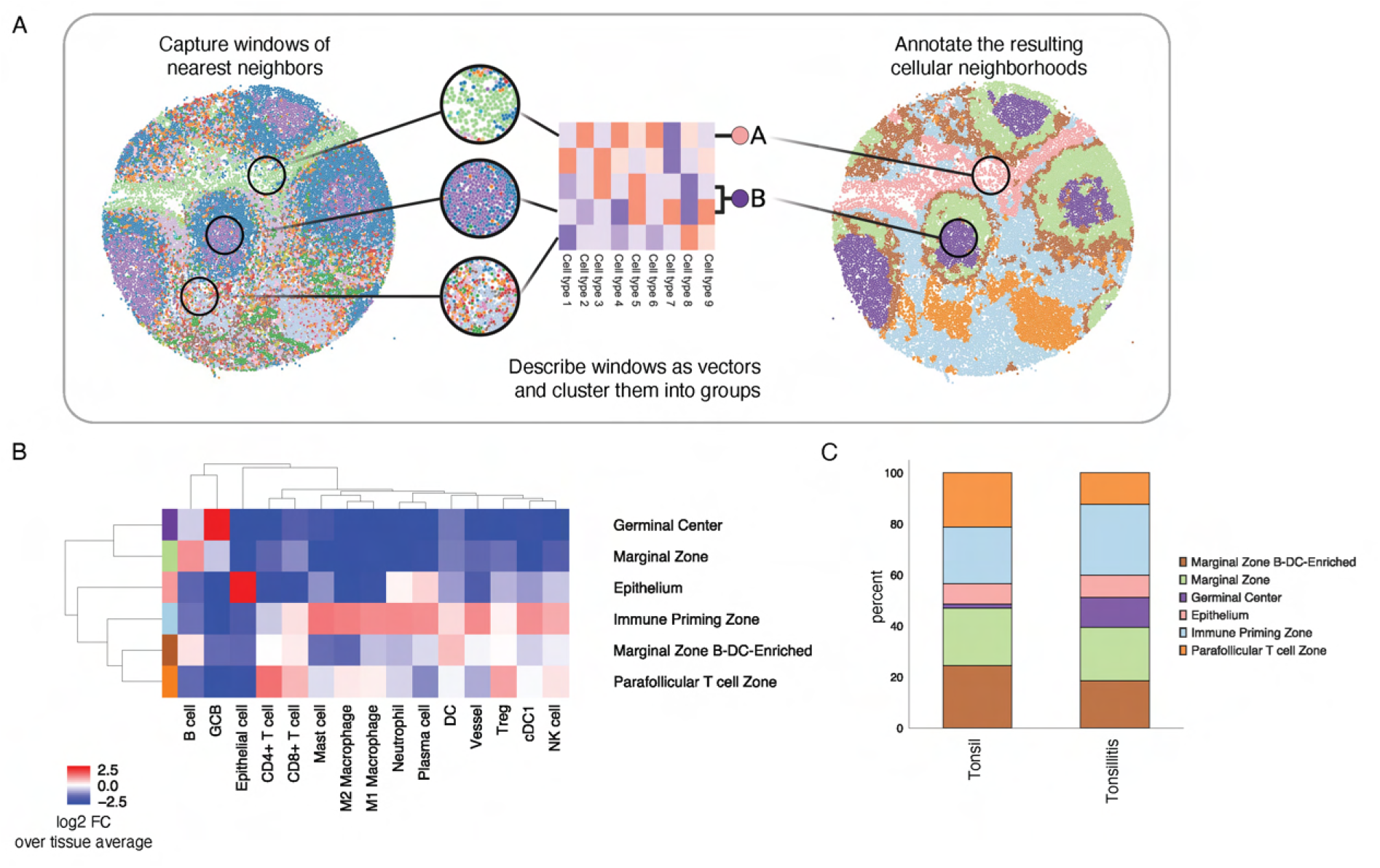
Cellular neighborhood analysis. **(A)** The computation process to identify CNs. A cell-type percentage vector is first computed for each sliding window. Then all cell-type percentage vectors are clustered. CNs are annotated manually based on marker expression. **(B)** In our study of CODEX data from healthy and inflamed tonsils, six distinct CNs were identified. The heatmap shows those cell types enriched in each neighborhood. **(C)** The annotated CNs mapped back onto original tissue coordinates in healthy (left) and inflamed (right) tonsils. Each dot represents a cell colored by its CN assignment.

##### F.3. Step 20: Generate spatial context map

Spatial context maps are created as previously described (32) and the mechanism behind them is described more in detail in the *Supplementary Notes*. A layered graph visualization can be used to understand the interactions between CNs in a hierarchical combination fashion. The spatial context maps build on the cellular neighborhood analysis concept, but instead of clustering neighboring vectors, the combination with the fewest CNs that include over 85% (backslash percent), or a userdefine percentage, of those CNs within the window is identified. This combination reveals prominent neighborhood associations that are termed spatial context features (**Supp. Fig. 5A**). Maps for healthy and inflamed tonsils are shown in **Fig. 2E**.

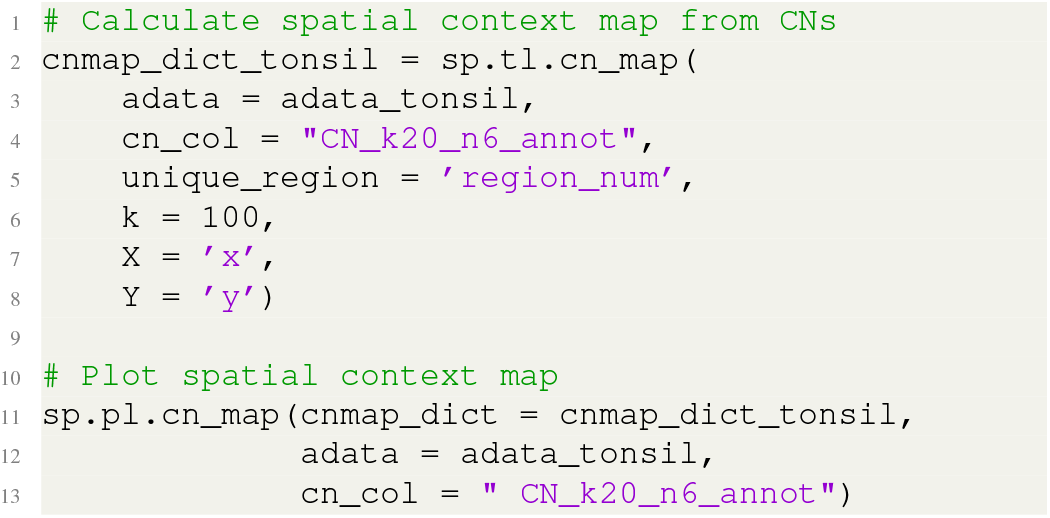

##### F.4. Step 21: Create the cellular neighborhood interface analysis via a barycentric coordinate plot

Biological interfaces between CNs are where specialized biological events take place (22). Users can visualize the CN interfaces via a barycentric coordinate system (**Supp. Fig. 6A**). In the barycentric coordinate system, each corner represents cells within the CN that are surrounded by cells assigned to the same neighborhood. Cells located on the edges of the triangle are indicative of the interface between two CNs, denoting windows with mixed compositions. Cells positioned at the center of the triangular barycentric coordinate system are situated in regions that have a mixture of three CNs. To zoom in on a region, three CNs of interest must be specified and a threshold of the minimum percentage of cells within a window that must belong to one of the three CNs for it to be displayed must be selected. A higher threshold results in purer representations of all three windows but fewer cells. In our example, we selected three CNs and inspected their interactions (**Fig. 2G**). We observed that in the inflamed tonsil, there are more cells interfacing between the germinal center CN and the marginal zone B cell- and dendritic cell-enriched CN than in the healthy tonsil.

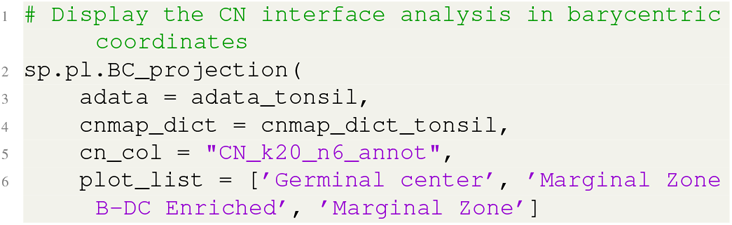
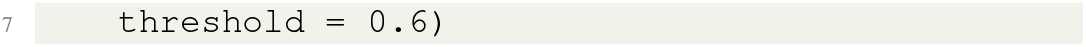

**Supplementary figure 5:**
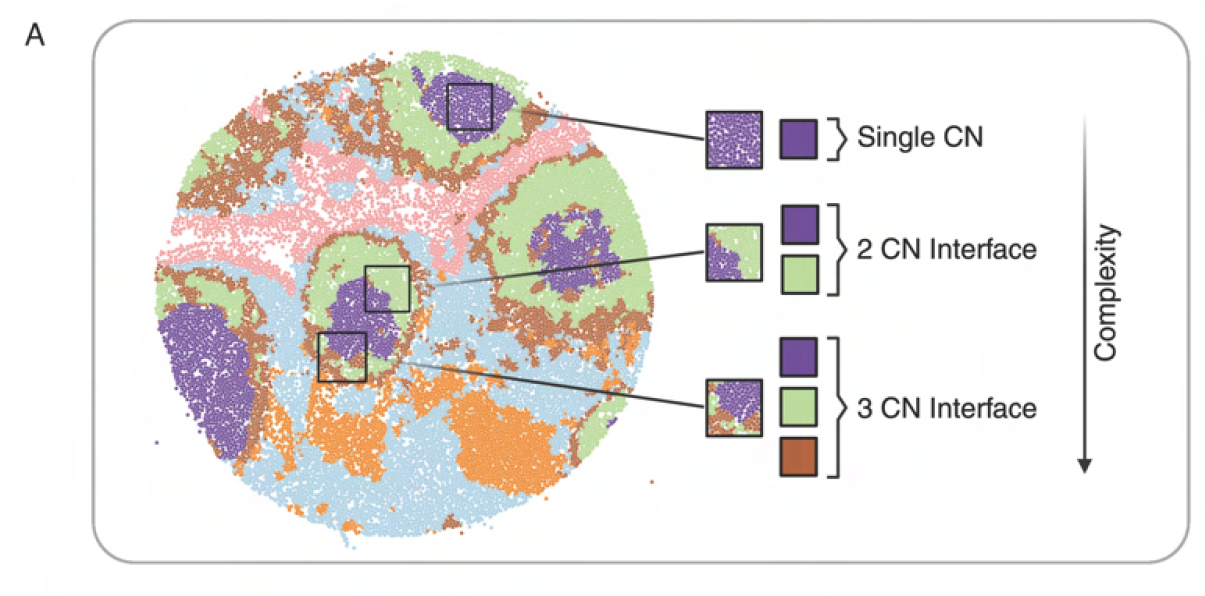
Spatial context map. **(A)** Schematic of spatial context map construction. Any combination of three user-defined CNs are analyzed. Relative proportional occurrences of these CNs are calculated and depicted on the barycentric coordinate coordinates. Rows show the number of neighborhoods in combinations: row 1, a single neighborhood accounts for at least 85% of the neighborhoods surrounding the window; row 2, two neighborhoods make up more than 85% of the neighborhoods in the window; and row 3 and beyond, multiple neighborhoods are present in the window.

**Supplementary figure 6:**
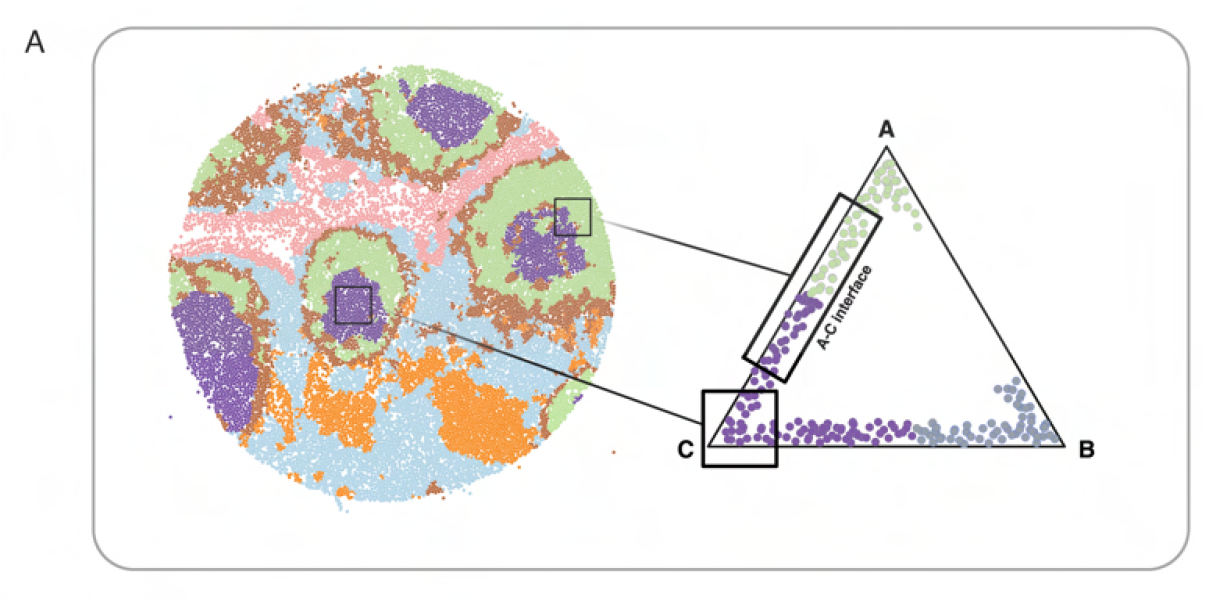
Cellular neighborhood interaction via barycentric coordinate plot. **(A)** Schematic of generation of barycentric plots of selected CN combinations. The initial selection criteria involve windows containing combinations of three CNs, each exceeding 85 percent of the total CN proportion. The relative frequencies of these selected neighborhoods are computed and visually represented in the barycentric coordinate system.

##### F.5. Step 22: Compute patch proximity

After inspection of the interfaces of different CNs, the compositions of the cells that comprise the interfaces. An analytic tool, patch proximity analysis (PPA), is created in SPACEc. PPA extracts continuous patches of a neighborhood using HDBSCAN clustering of the spatial coordinates within a region and then defines the cells that outline each patch (**Supp. Fig. 7A**). These cells are then used as anchor points to inspect cells within a user-defined radius (**Supp. Fig. 7B**). In our example data, we analyzed the cells surrounding the germinal center and observed that there were higher percentages of CD4+ T cells surrounding germinal center CNs in the tonsillitis sample than in the healthy tonsil sample (**Fig. 2H**).

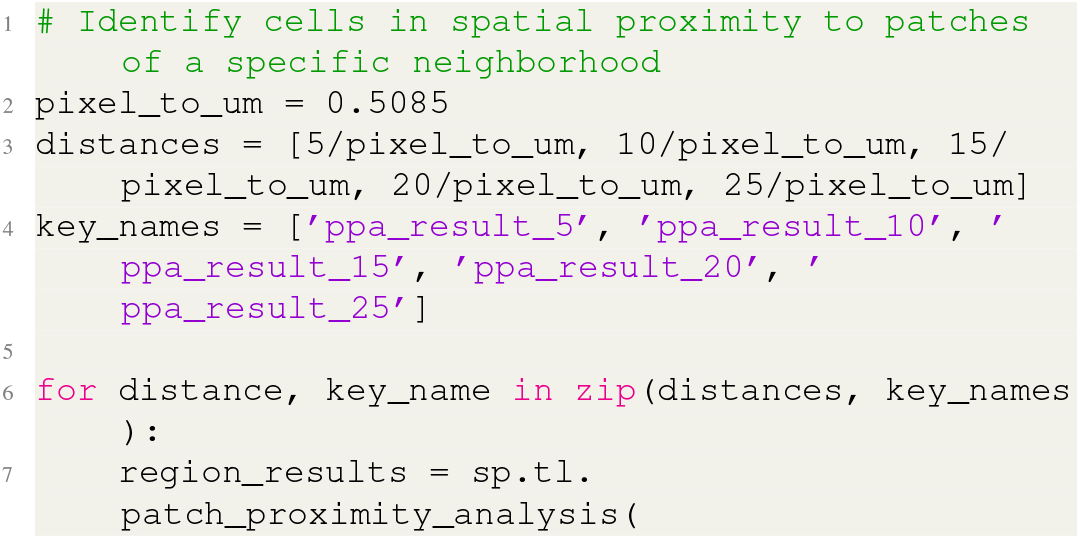
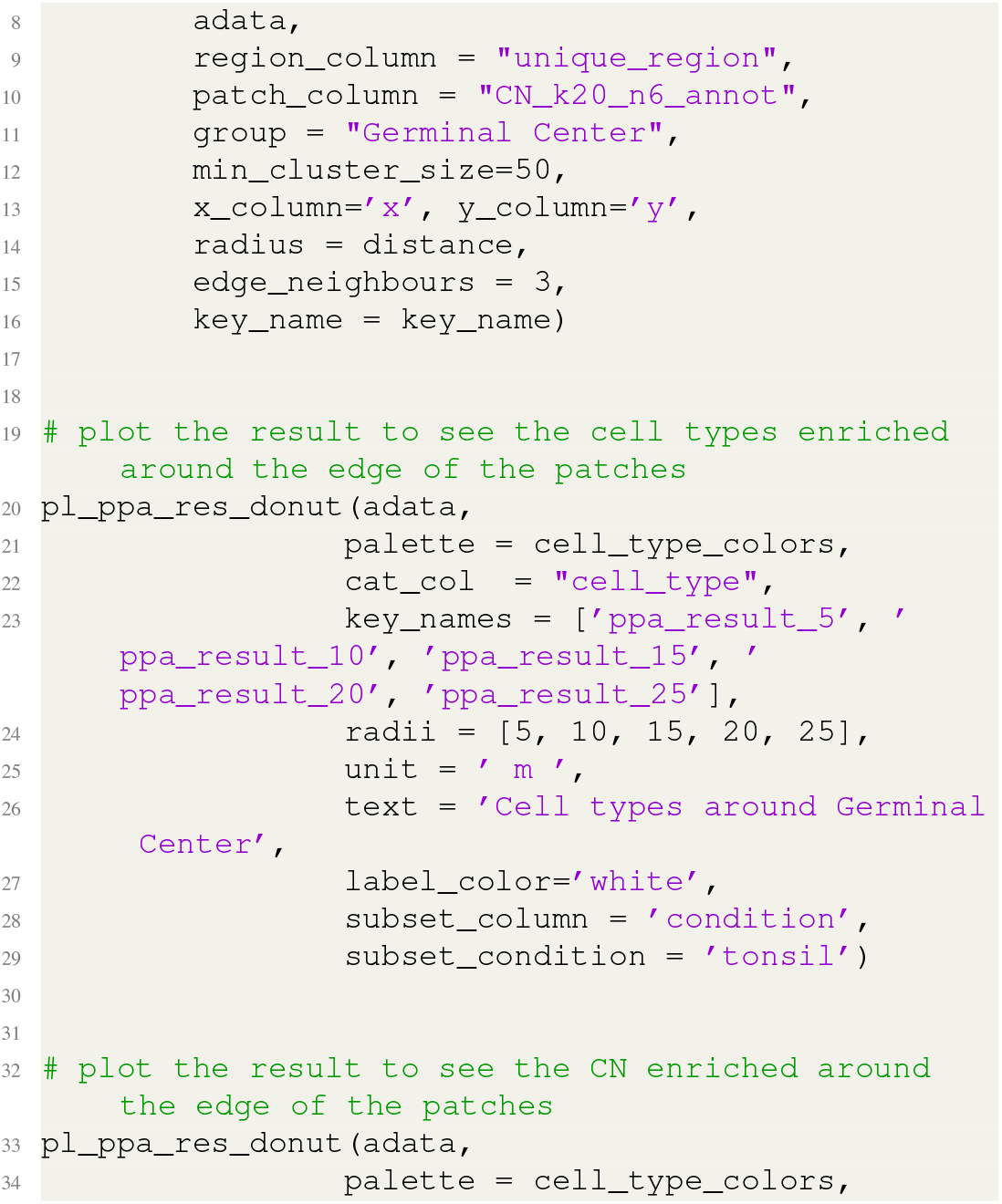
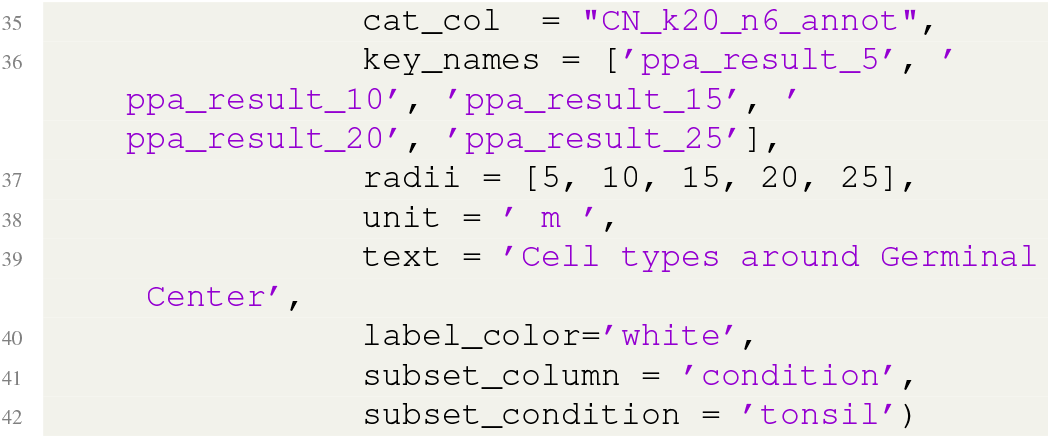

**Supplementary figure 7:**
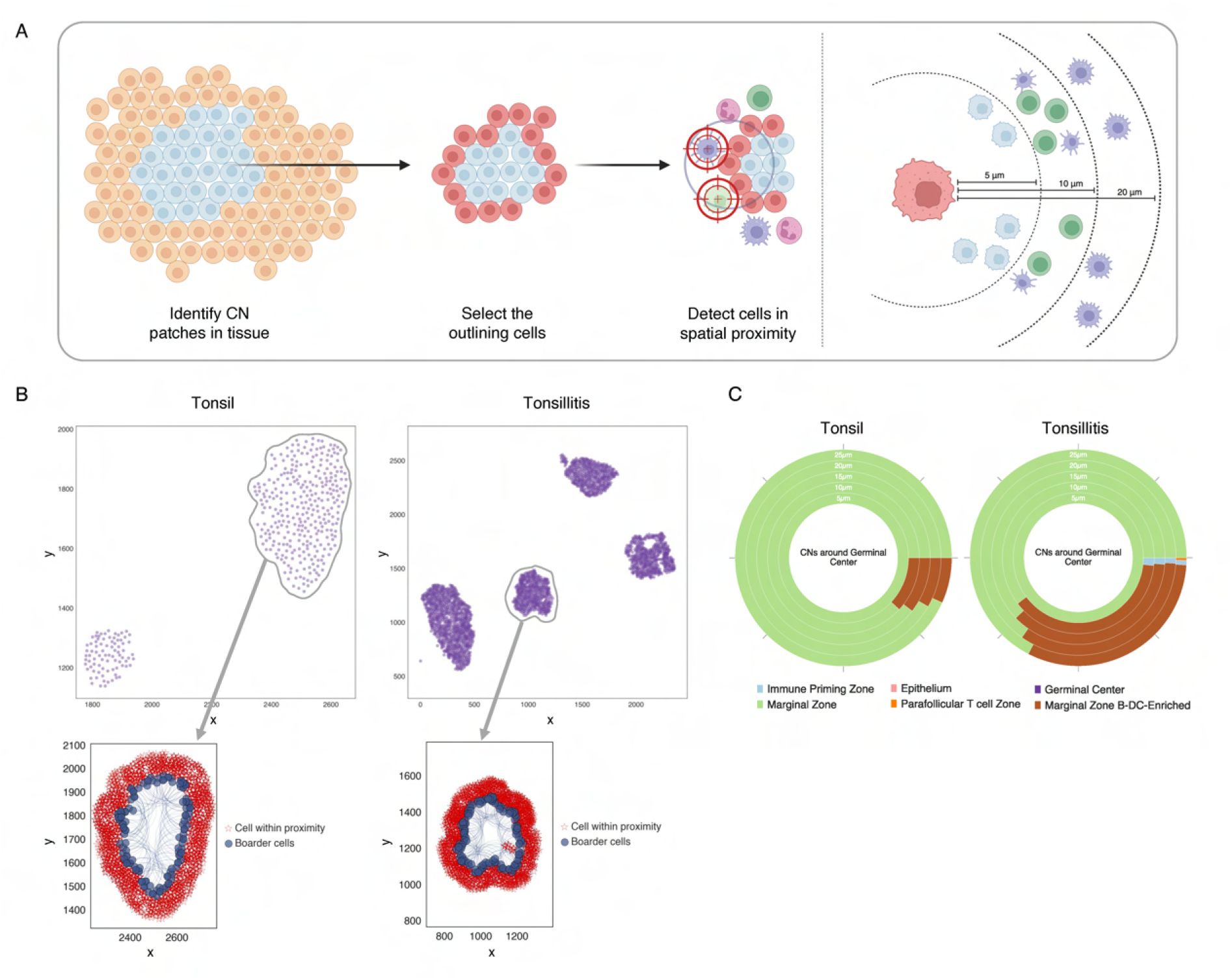
Patch proximity analysis. **(A)** Outline of the process of patch proximity analysis designed to identify cells surrounding each patch, which is defined by an isolated CN. Cells that border the patch within a user-defined radius are identified. **(B)** The surrounding cells within the user-defined proximity per each patch of the same CN are identified for the Germinal Center CN in the tonsil sample (left) and the tonsilitis sample (right). **(C)** Donut bar chart of percentages of cell types that surround all patches in the Germinal Center CN in healthy and inflamed tonsil samples. **(D)** Donut bar chart of percentages of CN that surround all patches in the Germinal Center CN in healthy and inflamed tonsil samples.

##### F.6. Step 23: Calculate cell-cell interactions

Cell-cell interaction analysis was described previously (21) and the algorithm is described in more detail in the *Supplementary Notes*. To understand how spatial distances between two cell types differ, the observed distances between all pairs of cell types within every region are measured and then the labels of the cell types are randomly shuffled to create a background distribution for comparison (**Supp. Fig. 8A**). In general, it is important to run large enough (e.g. 1000) iterations of the permutation per region to generate a null distribution. By contrasting the observed distances between cell-type pairs against the background distribution generated randomly, the probability of encountering such an occurrence relative to chance alone and the fold changes of the observed distance as compared to the expected distance can be calculated. The output of the algorithm contains the expected distances, observed distances, p-values, and log fold changes between each pair of cell types.

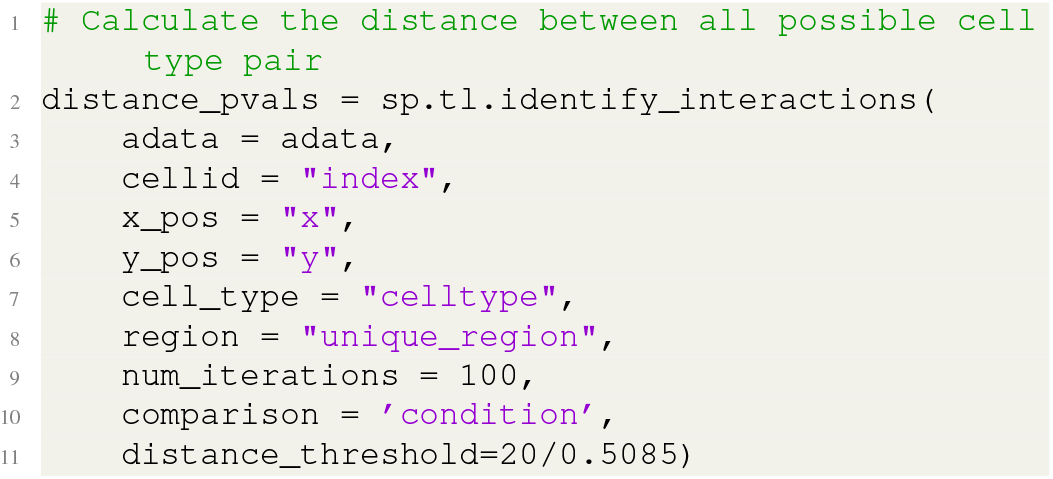

The output of the cell-cell interaction analysis is filtered to retain significant (p-value < 0.05 and fold changes > 0.1) interactions.

**Supplementary figure 8:**
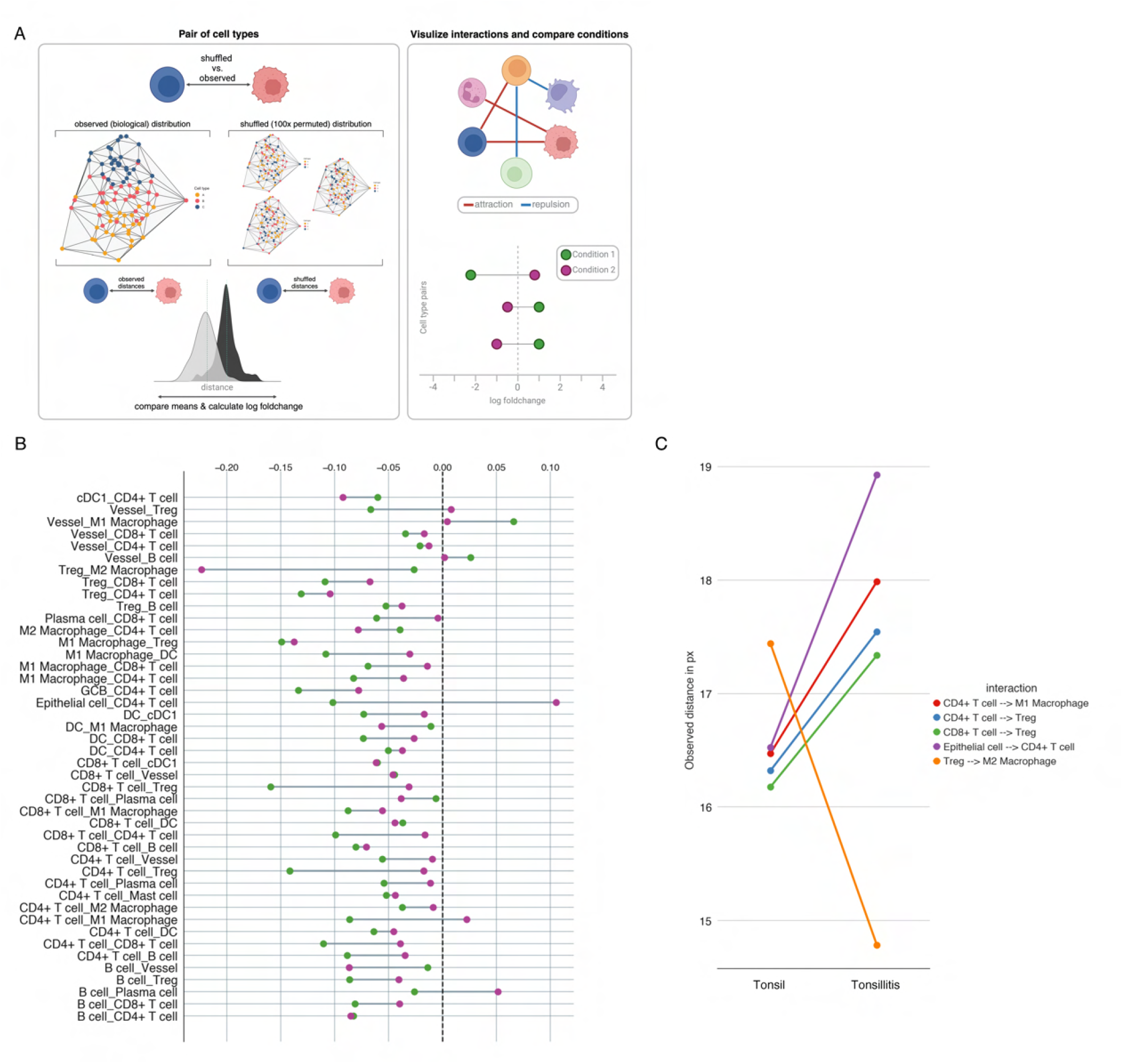
Cell-cell interaction analysis. **(A)** Left: Schematic of cell-cell interaction analysis. First, the original observed distances in the image are extracted. Next, labels for cell types are shuffled while maintaining fixed cell locations. This process is repeated x times as defined by the user. The aim is to generate a distribution of distances for each pair of cell types, simulating their observation under random chance conditions. By comparing the observed distances to the expected distances, meaningful statistics are derived. Right: Results can be visualized as distance network graphs and dumbbell plots. **(B)** Dumbbell plot of statistically significant differences in cell-type distances between the samples of healthy (green) and inflamed (purple) tonsil. **(C)** Bar chart of the five most significant interactions with the highest fold changes between healthy and inflamed tonsil.

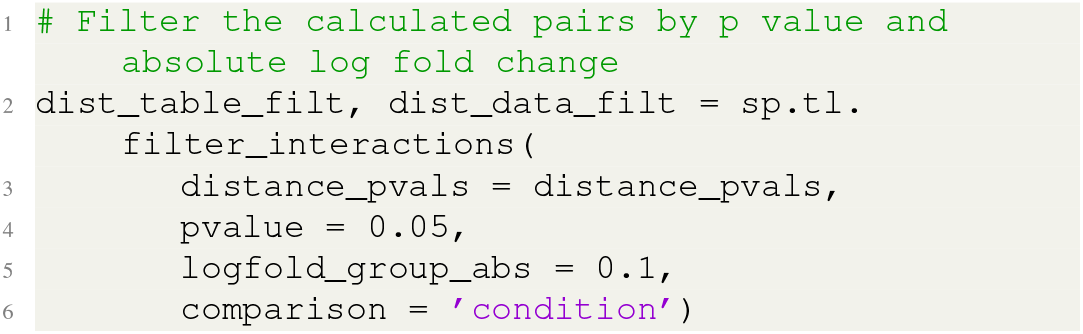

Multiple visualization possibilities are available within SPACEc (**Supp. Fig. 8B-C** and **Fig. 2I**).

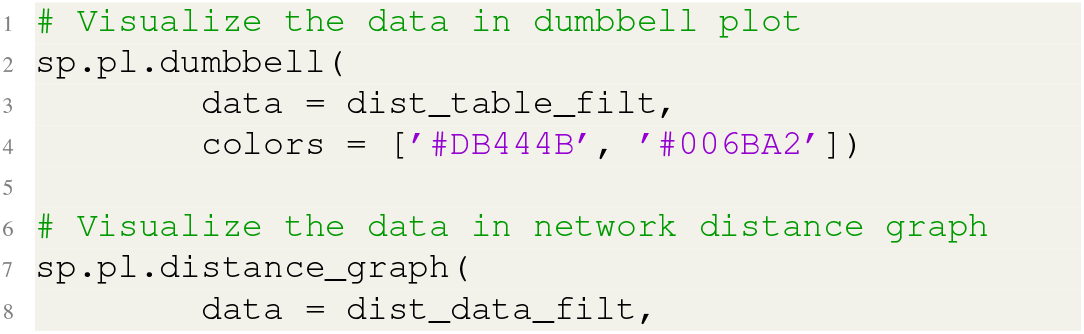
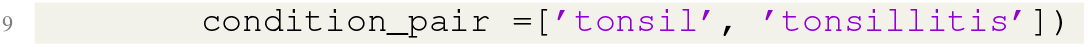

In our example data, we zoom in on the top 5 most significant cell-cell interactions and observed that the greatest significant distance changes between tonsil and tonsillitis cases are B cell to germinal center B cell, CD4 positive T cell to germinal center B cell, CD4 positive T cell to macrophage 1, dendrite cell to B cell, and Plasma cell to B cell (**Supp. Fig. 8C**).

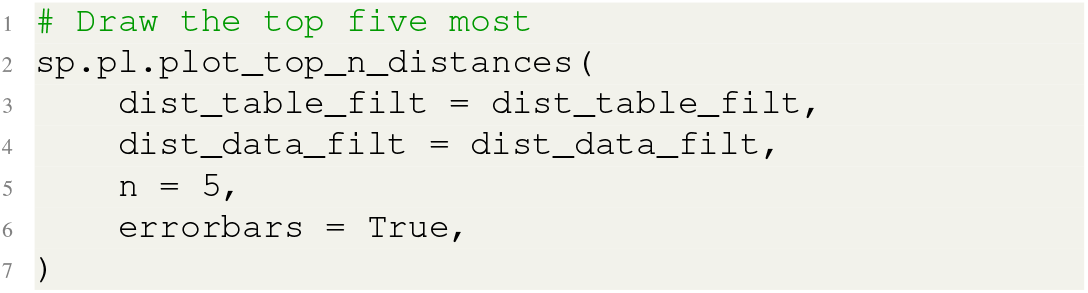

## Notes

### Competing Interest Statement

The authors have declared no competing interest.

